# The Generalized Haldane (GH) model tracking population size changes and resolving paradoxes of genetic drift

**DOI:** 10.1101/2024.02.19.581083

**Authors:** Yongsen Ruan, Xiaopei Wang, Mei Hou, Liying Huang, Wenjie Diao, Miles Tracy, Shuhua Xu, Weiwei Zhai, Zhongqi Liufu, Haijun Wen, Chung-I Wu

## Abstract

Population genetic models, such as the Wright-Fisher (WF) model, track relative gene frequencies. The absolute gene copy number, or population size (*N*), is supplied externally for tracking genetic drift. JBS Haldane (1927) proposed an alternative model based on the branching process, whereby each gene copy is transmitted to *K* descendants with the mean and variance of *E*(*K*) and *V*(*K*). In this model, *E*(*K*) governs *N*, while *V*(*K*)/*N* governs genetic drift. Nevertheless, as the branching process allows *N* to drift unboundedly, a Generalized Haldane (GH) model that regulates N more tightly is proposed. The GH model can account for several paradoxes of molecular evolution. Notably, genetic drift may often become stronger as *N* becomes larger in the ecological setting, thus contradicting the general view. In particular, a very small population growing exponentially experiences little drift. Interestingly, when the population grows and *N* oscillates near the carrying capacity, the paradoxical trend is also observed in both field works and laboratory experiments. This paradox whereby population size in genetics (*N*_*e*_) and ecology (*N*) could be negatively correlated is resolved by the GH model. Additional paradoxes include ii) The two sexes experiencing drift differently; iii) Genetic drift of advantageous mutations being independent of *N*; iv) Multi-copy gene systems (viruses, mitochondria, etc.) having no definable *N*_*e*_ (for effective *N*). In brief, the GH model defines genetic drift simply as *V*(*K*), or *V*(*K*)/*N* averaged over the population. It represents an attempt at integrating genetical and ecological analyses into one framework.

## Introduction

Genetic drift is defined as the random changes in frequency of genetic variants (Crow and Kimura 1970; Hartl and Clark 1997; Lynch, et al. 2016). Much, or even most, of DNA sequence evolution is governed by the random drift of neutral mutations. Nevertheless, even non-neutral variants are subjected to genetic drift as advantageous mutations often have a small probability of being fixed. If the strength of genetic drift is under-estimated, random changes that are missed in the analysis would result in the over-estimation of other evolutionary forces of greater biological interest, including selection, mutation, migration, meiotic drive and so on. In particular, the conclusion of pervasive selection at the molecular level has been suspected to be due to the failure to fully account for genetic drift (Crow and Kimura 1970; Fu 1997; Li 1997; Charlesworth 2009; Lynch, et al. 2016; Chen, Yang, et al. 2022).

Hagedoorn and Hagedoorn (1921) may be the first to suggest that random forces (accidents, for example) could impact long-term evolution. In the same decade, models of genetic drift were developed along two lines. The one approach that became widely adopted is the Wright-Fisher (WF) model (Crow and Kimura 1970; Hartl and Clark 1997), formalized by Fisher (1930) and Wright (1931). They define genetic drift to be due to the random sampling of genes. Thus, in generation transition, the variance in the frequency of a variant allele, *x*, would be

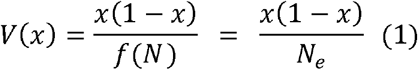

where *N* is the population size. (Here, we present the haploid model while a factor of 2 should be added to the diploid models.) The WF model has been modified in many directions (Wright 1969; Gillespie 1975; Eldon and Wakeley 2006; Chen, et al. 2017; Sackman, et al. 2019) whereby the effective population size, *N*_*e*_, is used to accommodate various deviations from the ideal WF population. Note that *N*, and hence *N*_*e*_, are imposed externally on the model as in the later Moran model (2008).

At about the same time, JBS Haldane introduced the branching process to model genetic drift (Haldane 1927; Haldane 1932). In the Haldane model, each gene copy is independently transmitted to *K* descendants with the mean and variance of *E*(*K*) and *V*(*K*). Again, we present the haploid model, noting some added complexities with diploidy.

The branching process of the Haldane model can generate *N* internally by formulating *N* of the next generation as *E*(*N′*) = *N*×*E*(*K*). Since Haldane’s model (1927) resets *N′* to a constant *N*, a Generalized Haldane (GH) model is proposed here that permits varying N under ecological regulation. Indeed, *E*(*K*) is often negatively correlated with *N* in actual populations (Smith and Slatkin 1973; Sibly and Hone 2002). The GH model thus tracks gene frequency changes as well as regulates *N*. In the Haldane model, *E*(*K*) would determine the trajectory of *N* changes and *V*(*K*) would determine the strength of genetic drift. Thus, there is no drift if *V*(*K*) = 0 irrespective of N.

The GH model is then applied to several paradoxes of genetic drift (see the overview below). It is possible that mathematical modifications of the conventional models (i.e., WF and Moran models) may resolve these paradoxes as well. Nevertheless, the key question is whether these modifications, often highly sophisticated, are biologically feasible (see Discussion). In comparison, the Haldane model is intuitively appealing and deserves to be considered as an alternative approach to genetic drift. As we study the ever more complex biological systems in the genomic era, such an alternative would seem desirable.

## Results

### Overview of the paradoxes of genetic drift

Among the paradoxes, a most curious one is genetic drift in relation to changing population size. The WF model dictates stronger drift when *N* is smaller. However, when *N* is very small (say, *N* = 10 in a bacteria or yeast population) and increases to 20, 40, 80 and so on, there is in fact little drift in this exponential phase. Drift will intensify when *N* grows to near the carrying capacity. This trend is the exact opposite of the standard view. The second paradox concerns the genetic drift of sex-linked genes in relation to sex-dependent breeding successes (Charlesworth and Charlesworth 2000; Bachtrog 2013; Cortez, et al. 2014; Wilson Sayres, et al. 2014; Makova, et al. 2024). A third paradox is about drift strength when selection is also at work. A new mutation with a selective advantage would always be fixed if there is no random drift. With drift, fixation is probabilistic, but the probability does not depend on *N* or *N*_*e*_. This curious property echoes the view that *N* is a scaling factor of drift that is determined by *V*(*K*) (see Results).

The power of the Haldane model in resolving these paradoxes is the focus of this study. Furthermore, the companion study (Wang et al. 2024) addresses a fourth paradox – the evolution of multi-copy gene systems, whereby evolution proceeds in two stages - between as well as within individuals. Multi-copy gene systems including viruses, mitochondria, transposons and ribosomal genes do not have easily definable Ne. Broadly speaking, even diploidy is also multi-copy systems (Silver 1985; Wu, et al. 1988; Wu, et al. 1989; Lindholm, et al. 2016; Courret, et al. 2019).

### On the Haldane model of genetic drift

In the original Haldane model, genetic drift is defined by the branching process (see Eq. (4) of Chen, et al. (2017)). Let *K* be the number of progenies receiving the gene copy of interest from a parent; *K* would follow the same distribution for all neutral variants. In haploids, *K* is equivalent to the progeny number of each parent. In diploids, *K* is the transmission success of each gene copy, which is separately tracked. By the Taylor approximations of the ratio of two variables (Kendall, et al. 2006), we obtain the approximation as

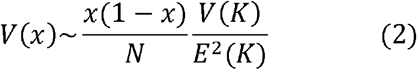

where *x* is the variant frequency (see Supplementary Information). This is the same approximation, when *V*(*K*) is not very large, as obtained by the WF model (Kimura and Crow 1964; see Chen et al. 2017 for details). For Eq. (2), we provide further extensive simulations on the accuracy in both the fixation probability and fixation time of neutral variants (see Supplementary Information). Note that N, constant or variable, is supplied externally to the WF model but it is tracked internally in the Haldane model (see the next section). The apparent equivalence of Eq. (2) makes a simple point that, between the WF and Haldane models, the mathematical results are often shared but the biological meanings may not be identical. (Similarly, the diffusion and coalescence theories are also distinct approaches to the same population genetic phenomena.)

In the Haldane model, there would be no genetic drift if *V*(*K*) = 0, regardless of the value of *N*. When *E*(*K*) = 1 and *N* stays constant, we obtain

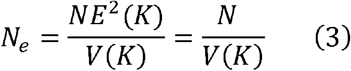

Eq. (2) and Eq. (3) have been obtained for the WF model (Kimura and Crow 1963; Crow and Denniston 1988; Chen, et al. 2017). The issue will be highly significant when sex-dependent drift is analyzed.

In **Table 1**, the data of progeny number from the literature indeed show over-dispersion in all taxa surveyed, i.e., *V*(*K*) > *E*(*K*). In fact, *V*(*K*) > 10 *E*(*K*) can often be found, e.g., the ratio of *V*(*K*)/*E*(*K*) among males of the mandrill (*Mandrillus sphinx*), the great reed warbler (*Acrocephalus arundinaceus*), and the rhesus macaque (*Macaca mulatta*) at 19.5, 12.6 and 11.3, respectively (Hasselquist 1995; Setchell, et al. 2005; Dubuc, et al. 2014). We have also found that *V*(*K*) may be far larger than *E*(*K*) in other biological systems such as viruses (Ruan, Luo, et al. 2021; Ruan, Wen, et al. 2021; Hou, et al. 2023).

**Table 1.** Field record of *E*(*K*) and *V*(*K*) across diverse taxas.

| Species | Type | Order | Gender | $E(K)$ | $V(K)$ | $V(K)/E(K)$ | Sample Size | Literature |
| --- | --- | --- | --- | --- | --- | --- | --- | --- |
| *Macaca mulatta* | Mammal | Primates | Female | 7.7 | 15.4 | 2.0 | 132 | Dubuc, et al. (2014) |
|  |  |  | Male | 8.7 | 98.7 | 11.3 | 86 |  |
| *Mandrillus sphinx* | Mammal | Primates | Female | 4.7 | 19.8 | 4.2 | 51 | Setchell, et al. (2005) |
|  |  |  | Male | 3.5 | 68.2 | 19.5 | 53 |  |
| *Macaca sylvanus* | Mammal | Primates | Female | 1.2 | 2.9 | 2.4 | 98 | Kuester, et al. (1995) |
|  |  |  | Male | 1.5 | 3.9 | 2.7 | 81 |  |
| *Peromyscus californicus* | Mammal | Rodentia | Female | 4.7 | 7.7 | 1.6 | 16 | Ribble (1992) |
|  |  |  | Male | 4.4 | 13.7 | 3.1 | 20 |  |
| *Turdus merula* | Bird | Passeriformes | Female | 4.9 | 23.4 | 4.8 | 80 | Wysocki, et al. (2019) |
|  |  |  | Male | 4.0 | 27.0 | 6.8 | 72 |  |
| *Acrocephalus arundinaceus* | Bird | Passeriformes | Female | 7.7 | 53.3 | 6.9 | 82 | Hasselquist (1995) |
|  |  |  | Male | 10.5 | 132.3 | 12.6 | 64 |  |
| *Charadrius nivosus* | Bird | Charadriiformes | Female | 1.7 | 5.3 | 3.1 | 105 | Herman and Colwell (2015) |
|  |  |  | Male | 2.2 | 9.0 | 4.1 | 90 |  |
| *Actitis macularia* | Bird | Charadriiformes | Female | 5.2 | 26.9 | 5.1 | 66 | Oring, et al. (1991) |
|  |  |  | Male | 3.3 | 17.2 | 5.2 | 103 |  |
| *Buteo buteo* | Bird | Accipitriformes | Female | 3.5 | 13.8 | 4.0 | 106 | Krüger and Lindström (2001) |
|  |  |  | Male | 2.7 | 9.3 | 3.4 | 132 |  |

### I. The first paradox – The paradox of changing *N*

#### 1. Empirical demonstration and simulation

This “paradox of changing *N*” is that “genetic drift increases when *N* increases” in direct contradiction with Eq. (2). **Fig. 1a** shows the results in a cell culture in the exponential growth phase where each cell doubles every 13 hours. Hence, *V*(*K*) ∼ 0 with almost no drift when *N* is as small as < 50. Genetic drift is expected to increase as *N* approaches the carrying capacity.

**Fig. 1.**
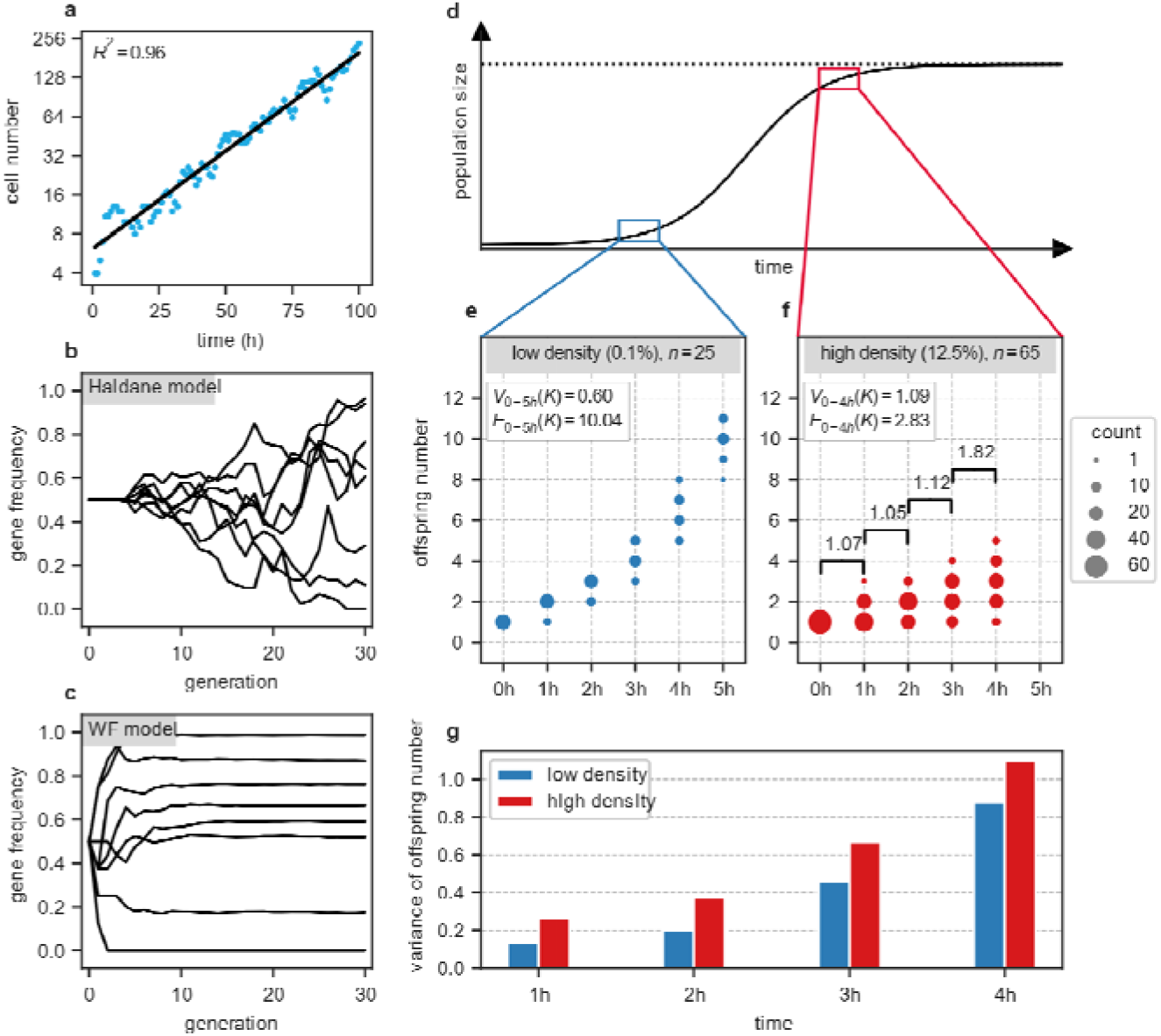
The paradox of genetic drift when the population size (*N*) changes. (**a**) In the laboratory cell culture, nearly all cells proliferate when *N* is very small (interpreted in the next panel). (**b**) The simulation of Haldane model shows little genetic drift in the exponential phase. As in (a), drift may increase due to the heightened competition as the population grows. (**c**) Simulation by the WF model shows a pattern of drift opposite of the Haldane model. (**d**) The patterns of drift at the low and high *N* are analyzed in the framework of the logistic growth model. (**e**-**f**) Measurements of genetic drift in laboratory yeast populations at low and high density as defined in (d). The progeny number of each cell, *K*, is counted over 4 or 5 intervals as shown by the dots, the sizes of which reflect the cell number. *E*(*K*) and *V*(*K*) are presented as well. In panel (**f**), the change of offspring number overtime, denoted as *V*(*ΔK*), is shown above the braces. (**g**) The variance of offspring number *V*(*K*) increases, observed in (e) and (f), as population size increases.

The paradox is exemplified through computer simulations in **Fig. 1b-c**. These panels show the drift pattern in the Haldane model and the standard WF model, respectively, when the populations are growing as the logistic growth model (see Methods). In the WF model, the drift is strong initially but weakened as *N* increases. The pattern is reversed in the Haldane model. The contrast between the two panels is the paradox.

To verify the simulations of **Fig. 1b-c**, we study the growth of yeast populations under laboratory conditions where cells appear to increase following the logistic growth curve. As shown in **Fig. 1d**, the lower portion of the S-curve portrays the exponential growth in a low-density population, while the upper curve indicates a slowdown in growth near the population’s carrying capacity. We directly recorded the individual output (*K*) of each yeast cell in the early (*n* = 25) and late (*n* = 65) stage of population growth under real-time high-content fluorescence imaging (Methods).

It’s evident that *V*(*K*) is decoupled from *E*(*K*). In the early stage, *E*(*K*) is 10.04 with *V*(*K*) = 0.60 over five one-hour intervals. In the high-density phase, *E*(*K*) decreases to 2.83 but *V*(*K*) increases to 1.09 in four intervals (**Fig. 1e-f**). Most interestingly, during the late stage of population growth, *V*(*ΔK*) (adjusted to the four-hour time span for comparison; see Methods) indeed increases as *N* increases (**Supplementary Table 3**). At the last time point, the population is closest to the carrying capacity and *V*(*ΔK*) experiences a substantial leap from around 1.12 to 1.82. Overall, the variance in progeny number *V*(*K*) in a high-density population is always greater that in a low-density population (**Fig. 1g**). Indeed, *V*(*K*) may exhibit an increase with the increase in *N*, sometimes outpaces the latter.

Measurements of these two growth phases demonstrate the “paradox of changing *N*” by showing i) there is almost no drift when *N* is very small; ii) as *N* increases, *V*(*K*), hence, genetic drift would also increase. Furthermore, *V*(*K*) may increase sharply when *N* is close to the carrying capacity. This trend appears to echo the field observations by Coltman, et al. (1999) who show that *V*(*K*) increases more than 5-fold when *N* increases only 2.5-fold in the United Kingdom (UK) population of Soay rams (*Ovis aries*).

#### 2. The density-dependent Haldane (DDH) model – A first attempt at the Generalized Haldane model

To explain the paradox of changing *N*, the model has to track and regulate *N*. Ironically, although the branching process can track the changes, the process has been deemed unsuitable for population genetics. This is because *N*_*t*_ = *N*_0_ *E*(*K*)^*t* would drive *N* to approach either zero or infinity as *t* becomes very large. Indeed, Haldane’s original model reset *N′* to *N* in each generation. Any generalized Haldane (GH) model must take advantage of this feature by regulating *N* via *E*(*K*), as attempted in this section.

The “paradox of changing *N*” can be explained by *N*_*e*_ = *N*/*V*(*K*) if the numerator and denominator move in the same direction. Most interestingly, *V*(*K*) may increase more than *N* itself. This has been shown when *V*(*K*) is close to zero while *N* is growing exponentially. In addition, when *N* is very near the carrying capacity, even a small increase in *N* would intensify the competition and inflate *V*(*K*). This is shown in the yeast experiment of **Fig. 1d** and supported by Coltman, et al. (1999) field study of ram populations in UK. Thus, the relationship between *N* and *V*(*K*) may realistically lead to this paradox.

We now extend the Haldane model by incorporating density-dependent regulation of *N* in order to define the conditions of this paradox. The model developed here is on an ecological, rather than an evolutionary time scale. The simplest model of *N* regulation is the logistic equation:

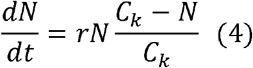

with *C*_*k*_ being the carrying capacity. In the ecological time scale, a changing *N* would mean that the population is departing from *C*_*k*_ or moving toward it (**Fig. 2a**). (As will be addressed in Discussion, changing *N* in the WF model is depicted in **Fig. 2b** whereby *N* is at *C*_*k*_. *C*_*k*_ may evolve too, albeit much more slowly in an evolutionary time scale.) Here, we consider changes in *N* in the ecological time scale. Examples include the exponential growth when *N* ≪ *C*_*k*_, seasonal fluctuation in *N* (Frankham 1995; Charlesworth 2009) and competition modeled by the Lotka-Volterra equations (Bomze 1983).

**Fig. 2.**
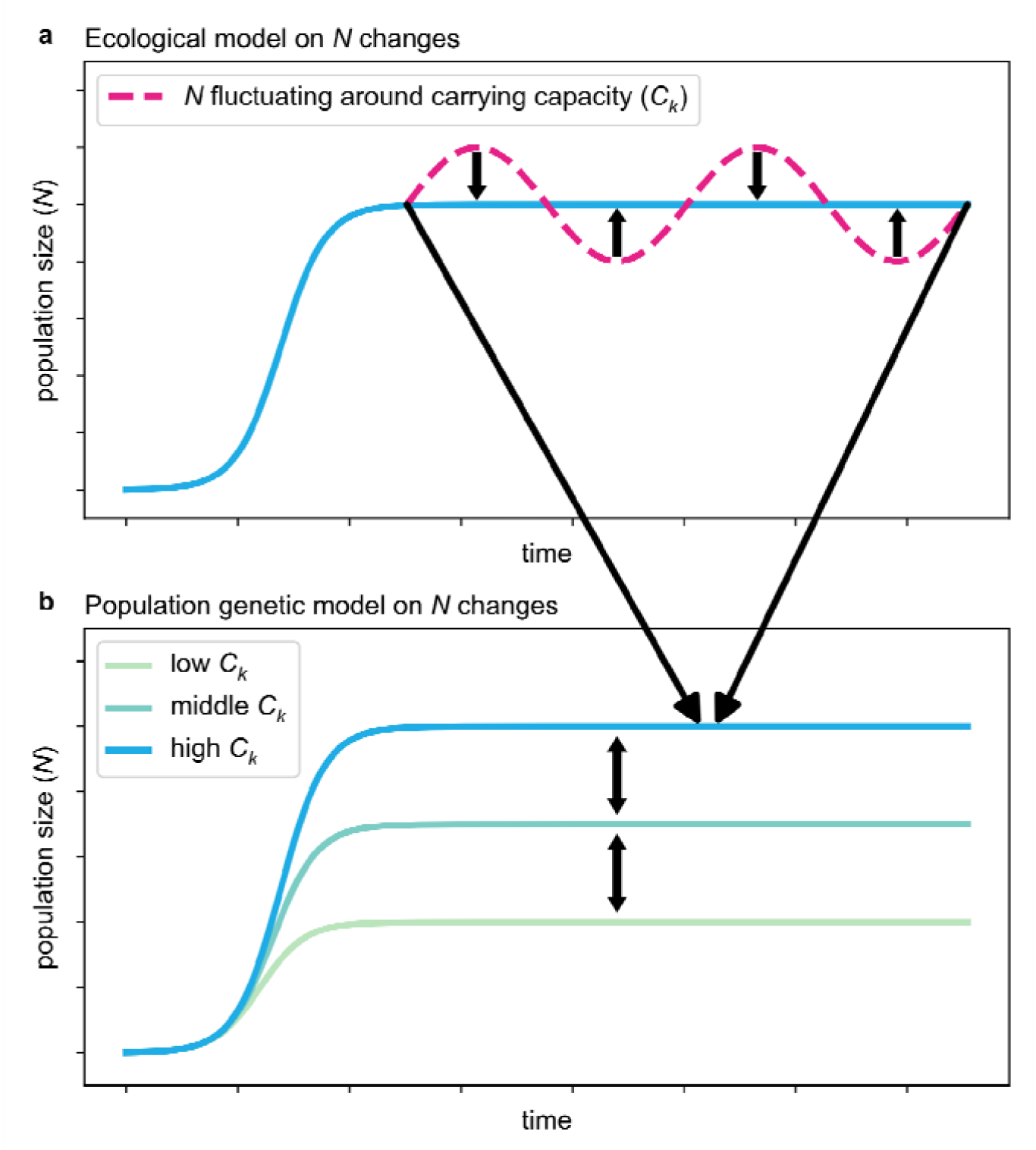
The meaning of population size (*N*) changes in ecology vs. in population genetics. (**a**) In ecology, changing *N* would generally mean a population approaching or departing the carrying capacity. (**b**) In population genetics, a population of size *N* is assumed to be at the carrying capacity, *C*_*k*_. Thus, changes in *N* would mean an evolving *C*_*k*_, likely the consequence of environmental changes. The arrows indicate the disparity in time scale between the two scenarios.

Details of the DDH model are given in the Supplementary Information. A synopsis is given here: We consider a non-overlapping haploid population with two neutral alleles. The population size at time *t* is *N*_*t*_. We assume that expected growth rate *E*(*K*) is greater than 1 when *N* < *C*_*k*_ and less than 1 when *N* > *C*_*k*_, as defined by Eq. (5) below:

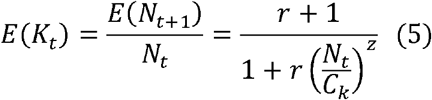

The slope of *E*(*K*) vs. *N* (i.e., the sensitive of growth rate to changes in population size), as shown in **Fig 3a**, depends on *z*. To determine the variance *V*(*K*), we assume that *K* follows the negative binomial distribution whereby parents would suffer reproduction-arresting injury with a probability of *p*_*t*_ at each birthing (Supplementary Information). Accordingly, *V*(*K*) can then be expressed as

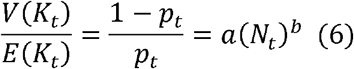

**Fig. 3.**
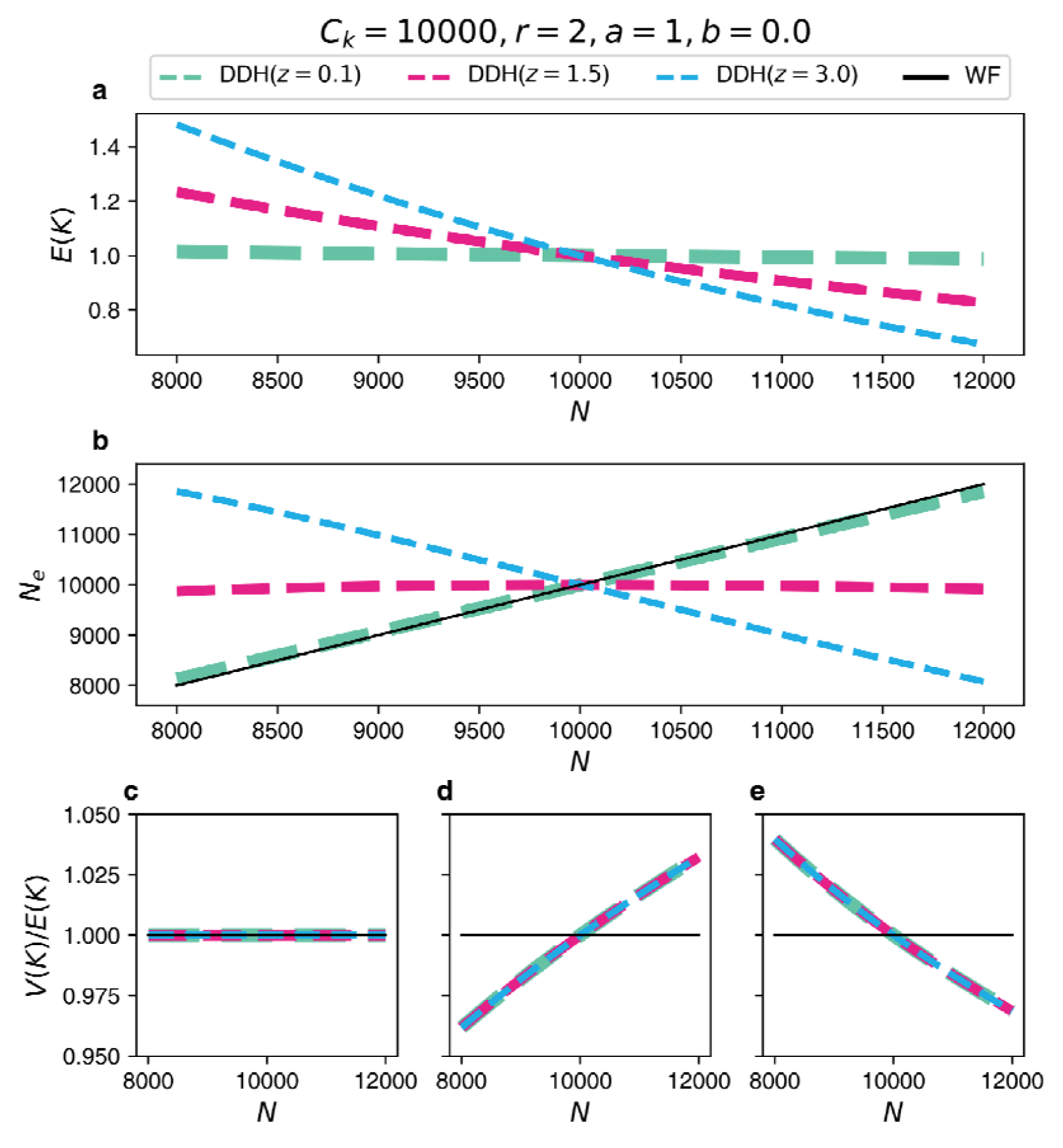
Genetic drift as a function of population size in the DDH model. For all panels, the carrying capacity is *C*_*k*_ = 10,000 and the intrinsic growth rate is *r* = 2. (**a**) When *N* increases, *E*(*K*) decreases as modeled in Eq. (5). The *z* value of Eq. (5) (0.1, 1.5 and 3) determines the strength of *N* regulation, indicated by the slope of *E*(*K*) near *C*_*k*_ = 10,000. (**b**) Depending on the strength of *N* regulation near *C*_*k*_, genetic drift can indeed decrease, increase or stay nearly constant as the population size increases. Thus, the conventional view of *N*_*e*_ being positively dependent on *N* is true only when the regulation of *N* is weak (the green line). At an intermediate strength (the red line), *N*_*e*_ is nearly independent of *N*. When the regulation becomes even stronger at z = 3, *N*_*e*_ becomes negatively dependent on *N*. (**c-e**) *V*(*K*)/*E*(*K*) of Eq. (6)) is shown as a function of *N*. The results of panel (b) are based on a constant *V*(*K*)/*E*(*K*) shown in panel (c). Interestingly, the results of panel (b) would not be perceptibly changed when *V*(*K*)/*E*(*K*) varies, as shown in panel (d) and (e).

By Eq. (6), the ratio of *V*(*K*)/*E*(*K*) could be constant, decrease or increase with the increase of population size. With *E*(*K*) and *V*(*K*) defined, we could obtain the effective population size by substituting Eq. (5) and Eq. (6) into Eq. (3).

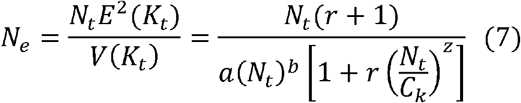

Eq. (7) presents the relationship between effective population size (*N*_*e*_) and the population size (*N*) as shown in **Fig. 3**. The density-dependent *E*(*K*) could regulate *N* with different strength (**Fig. 3a**). The steeper the slope in **Fig. 3a**, the stronger the regulation.

The main results of this study are depicted in **Fig. 3b**. First, with no or little regulation of *N, N*_*e*_ and *N* are strongly correlated. The green dashed lines portray the conventional view of decreasing drift with increasing *N*. Second, under a particular strength of *N* regulation, the red line shows no dependence of *N*_*e*_ on *N*, meaning that genetic drift is independent of *N*. Finally, as *N* becomes strongly regulated, *N*_*e*_ and *N* would be negatively correlated as the blue dashed line shows. This trend is the paradox of changing *N*.

In summary, genetic drift effect can indeed decrease, increase or stay nearly constant as the population size increases, depending on the strength of *N* regulation near *C*_*k*_. We further note that the *V*(*K*)/*E*(*K*) ratio of Eq. (7) can be independent (**Fig. 3c**), positively dependent (**Fig. 3d**), or negatively dependent (**Fig. 3e**) on *N*. Interestingly, the results of **Fig. 3b** are nearly identical when the *V*(*K*)/*E*(*K*) ratio changes (**Supplementary Figs 1-3**). These results show that the strength of genetic drift depends on the ecology that governs *E*(*K*), *V*(*K*) and *N*.

The paradox of changing *N* may not be an exception but a common rule in natural populations. Note that the WF approximation yielding Eq. (2) assumes nearly constant *N*. The assumption would imply very strong *N* regulation near *C*_*k*_, which is precisely the condition leading to the paradox of changing *N* (**Fig. 3**). Coltman et al. (1999)’s observations of the reproduction in rams is such a case.

### II. The paradox of genetic drift in sex chromosomes

Since the relative numbers of Y, X and each autosome (A) in the human population are 1:3:4, the WF model would predict the genetic diversity (θ_*Y*_, θ_*X*_ and θ_*A*_, respectively) to be proportional to 1:3:4. In a survey of human and primate genetic diversity, *θ*_*Y*_ is almost always less than expected as has been commonly reported (Hammer, et al. 2001; Wang, et al. 2014; Wilson Sayres, et al. 2014; Makova, et al. 2024). The low *θ*_*Y*_ value has been used to suggest that human Y chromosomes are under either positive or purifying selection (Wang, et al. 2014; Wilson Sayres, et al. 2014; Makova, et al. 2024). Similarly, the reduced diversity X-linked genes was interpreted as a signature of positive selection (Pan, Liu, et al. 2022).

As pointed out above, under-estimation of genetic drift is a major cause of over-estimation of selective strength. Our goal is hence to see if genetic drift of different strength between sexes can account for the observed genetic diversities. Details will be presented in Wang et al. (2024). Below is a synopsis.

Let *V*_*m*_ and *V*_*f*_ be the *V*(*K*) for males and females respectively and *α*′ = *V*_*m*_/*V*_*f*_ is our focus. (We use *α*′ for the male-to-female ratio in *V*(*K*) since *α* is commonly used for the ratio of mutation rate (Miyata, et al. 1987; Makova and Li 2002).) The three ratios, *θ*_*Y*_ /*θ*_*X*_ (denoted as *R*_*YX*_), *θ*_*Y*_ /*θ*_*A*_ (*R*_*YA*_) and *θ*_*X*_ /*θ*_*A*_ (*R*_*XA*_), could be expressed as the functions of *α*′, which incorporate the relative mutation rates of autosomes, X and Y (Supplementary Information). These relative mutation rates can be obtained by interspecific comparisons as done by Makova and Li (2002).

We note that there are many measures for the within-species genetic diversity, *θ* = 4*N*_*e*_*μ*. Under strict neutrality and demographic equilibrium, these measures should all converge. In the neutral equilibrium, the infinite site model dictates the frequency spectrum to be *ξ*_*i*_ = *θ*/*i*, where *ξ*_*i*_ is the number of sites with the variant occurring *i* times in *n* samples. Since every frequency bin is a measure of *θ*, different measures put different weights on the *i*-th bin (Fay and Wu 2000; Fu 2022). While *π*, the mean pairwise differences between sequences, is most commonly used in the literature, we use several statistics to minimize the possible influences of selection and demography (Wang et al. 2024). In this synopsis, we used the Watterson estimator (Watterson 1975) as the measure for *θ*_*Y*_, *θ*_*X*_ and *θ*_*A*_ by counting the number of segregating sites.

Using any of the three ratios (*θ*_*Y*_ /*θ*_*X*_, *θ*_*Y*_ /*θ*_*A*_ and *θ*_*X*_ /*θ*_*A*_), we can obtain *α*′; for example, *R*_*YA*_ = *θ*_*Y*_/*θ*_*A*_ and, hence,

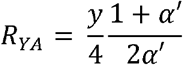

where *y* is the mutation rate of Y-linked sequences relative to autosomal sequences. With rearrangement,

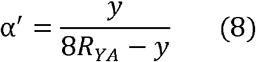

Similar formulae of *α*′ can be obtained for *R*_*YX*_ and *R*_*XA*_ but the accuracy for estimating *α*′ is highest by *R*_*YA*_ whereas *R*_*XA*_ is the least accurate for this purpose (Supplementary Information).

**Table 2** presents the *α*′ estimates in chimpanzees and bonobos. This is part of a general survey in mammals with a strong emphasis on primates and humans (Wang et al. 2024). It is almost always true that *α*′ > 5 in primates. Sources of data used are also given in **Table 2**. As shown for chimpanzees, *α*′ is often far larger than 5, above which the resolution is very low as can be seen in Eq. (8). Note that, when *R*_*YA*_ is under-estimated and approaching *y*/8, *α*′ would increase rapidly when the denominator is close to 0. That is why *α*′ often becomes infinity (**Supplementary Fig. 5**). The estimated *α*′ (MSE) in **Table 2** alleviates the problem somewhat by using all three ratios to calculate the mean square error (MSE) (Supplementary Information).

**Table 2.**
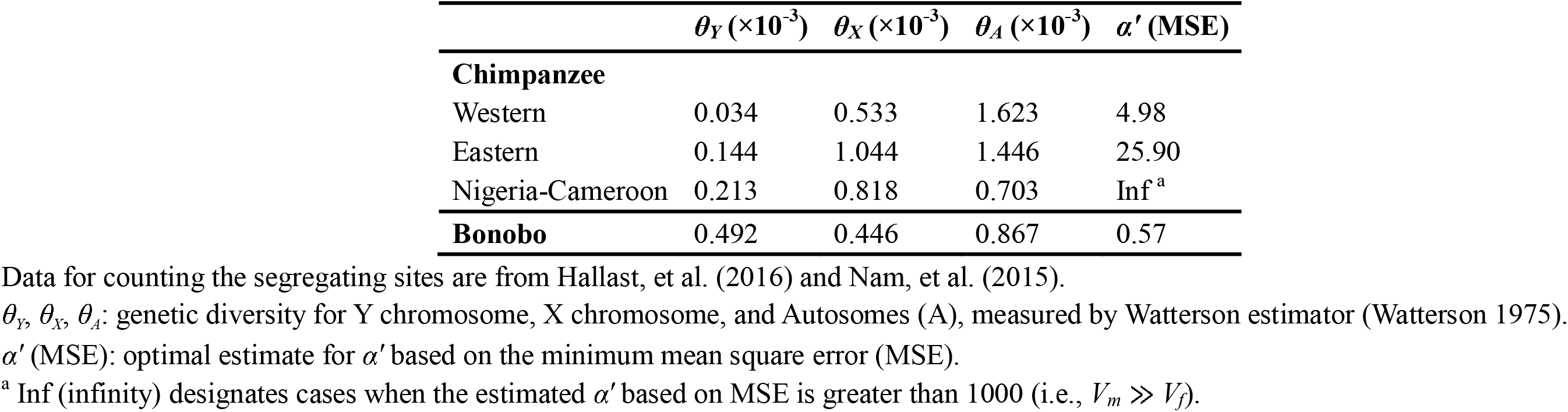
Estimation of *V*_*m*_/*V*_*f*_ in chimpanzee and bonobo.

Among primates surveyed, bonobo is the only exception with *α*′ < 1. While chimpanzees and bonobos are each’s closest relatives, their sexual behaviors are very divergent (de Waal 1995; De Waal and Lanting 2023). With unusually strong matriarchs, bonobo society seems to stand out among primates in sexual dominance. The *α*′ value is important in behavioral studies and, particularly, in primatology and hominoid research. If 0.1% of males sire 100 children and the rest have the same *K* distribution as females, *V*_*m*_/*V*_*f*_ would be ∼ 10. Such outlier contribution is the equivalent of super-spreaders in viral evolution which may easily be missed in field studies. In this brief exposition, we highlight again that *V*(*K*) or, more specifically, *V*_*m*_ and *V*_*f*_ are key to genetic drift. The differences in drift strength among X, Y and autosomes constitute a curious (but not immediately recognizable) paradox for the WF models as will be addressed in Discussion.

### III. The paradox of genetic drift under selection

Genetic drift operates on neutral as well as non-neutral mutations. Let us assume a new mutation, M, with a frequency of 1/*N* is fixed in the population with the probability of *P*_*f*_. Fisher (1930) first suggested that the fixation probability of an advantageous mutation should increase in growing populations, while decrease in shrinking populations. If there is no genetic drift, a beneficial mutation will always be fixed and *P*_*f*_ = 1. In the WF model, it is well known that *P*_*f*_ ∼ 2*s* for a new advantageous mutation, with fitness gain of *s*, while population size is large (Haldane 1932; Kimura 1962). This seems paradoxical that the determinant of genetic drift, 1/*N*_*e*_, does not influence *P*_*f*_ (Otto and Whitlock 1997; Lanfear, et al. 2014).

In the Method section, we show that *P*_*f*_ under the Haldane model is given by

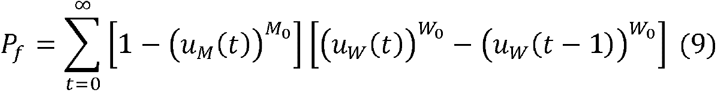

where *M*_0_ and *W*_0_ are the initial number of mutant allele (M allele) and wildtype allele (W allele), with *N* = *M*_0_ + *W*_0_. We note that *u*_*M*_(*t*) and *u*_*W*_(*t*), respectively, are the extinction probabilities of M allele and W allele by generation *t* with the initial number of alleles of 1. While obtaining the direct analytical solution of Eq. (9) may not be feasible, we could obtain its numerical solution due to its convergence as *t* increases (see Methods). The accuracy of the numerical solution from Eq. (9) is confirmed through simulation (**Supplementary Fig. 4**). Moreover, with the aid of the WF model (see Supplementary Information), we could obtain the approximation of Eq. (9) as follows.

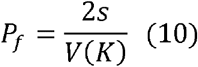

We verify Eq. (10) by both the numerical solution from Eq. (9) and simulations based on the branching process (**Supplementary Fig. 4**). The fixation probabilities obtained by numerical solution vs. those inferred from Eq. (10) are shown in **Fig. 4**. The salient feature is that the fixation probability of 2*s*, as in the classical formula, would be a substantial over-estimate when *V*(*K*) is larger than *E*(*K*). Eq. (10) is sufficiently accurate as long as *N*≥50. When *N* is as small as 10, the theoretical result is biased. Indeed, at such a low *N* value, the population is prone to extinction. The DDH model presented above should rectify this deficiency.

**Fig. 4.**
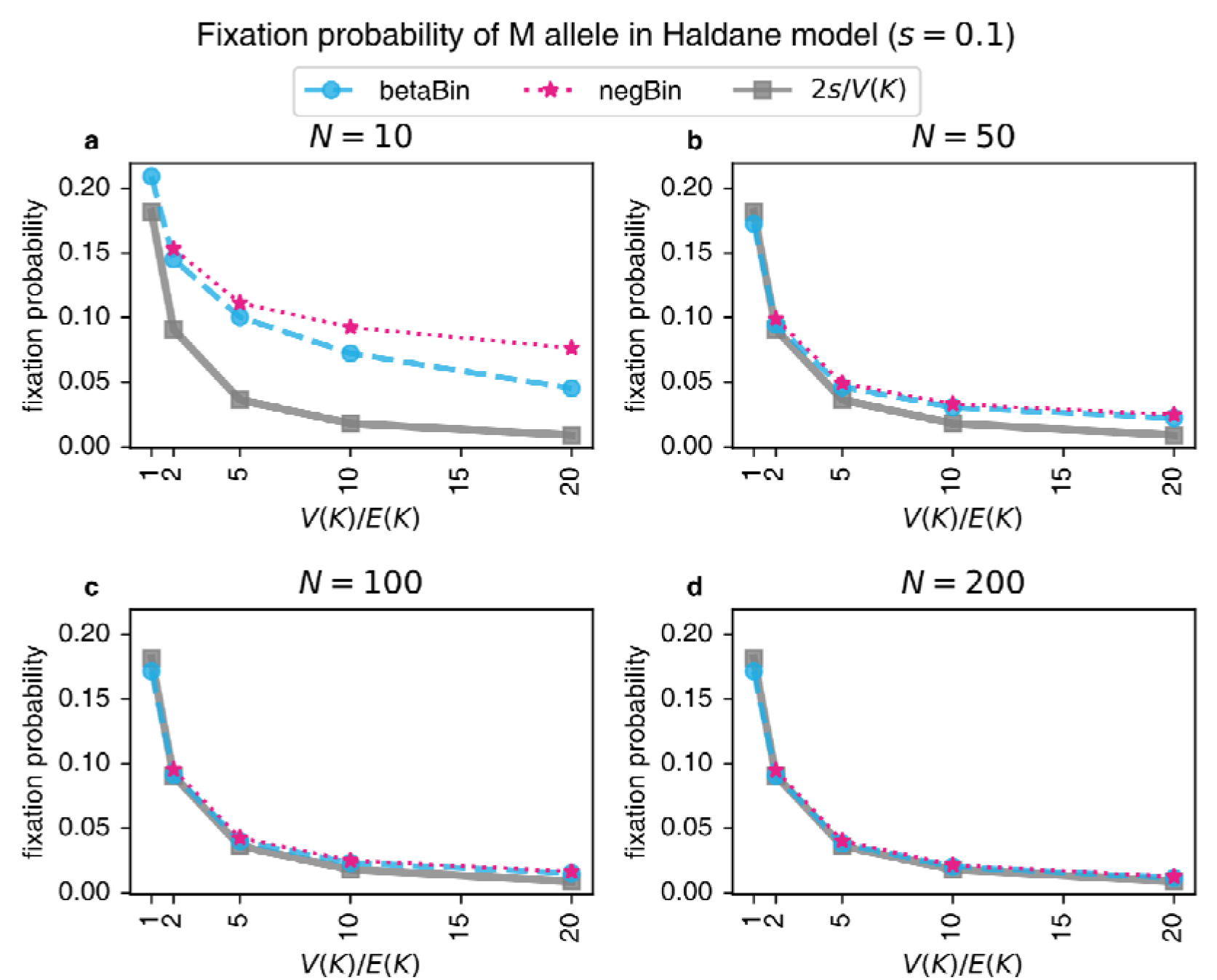
Fixation probability of a new advantageous mutation in the Haldane model. The fixation probabilities of a new advantageous mutation with the selective advantage of *s* = 0.1 are calculated based on approximate solution from Eq. (9) (i.e., 2*s*/*V*(*K*)) as well as numerical solution from Eq. (10). The numerical solution from Eq. (10) has been confirmed accurate by simulations (Supplementary Fig. 4). **(a-b)** When *N* < 50, the approximate fixation probability (the gray line) is lower than the simulated values (the color lines) due to population extinction. **(c-d)** By the Haldane model, the expected fixation probability of Eq. (9) is accurate when *N* reaches 100, as in most natural populations.

The main message of Eq. (10) is that genetic drift under positive selection is influenced by *s* and *V*(*K*) but not by *N*. This independence from *N* is explicable: when an advantageous mutation increases to a certain level still far below *N* (depending mainly on *s*), its fixation would be almost certain. This may be a most direct argument against equating genetic drift with *N*, or *N*_*e*_ (which is a function of changing *N*’s).

## Discussion

Genetic drift should broadly include all random forces affecting evolutionary trajectories. Hence, when *N* is not regulated by the models themselves but supplied externally, some random forces are excluded from the modeling. The DDH model may be a first-generation GH model that incorporates *N* regulation. We note that the original Haldane model suppresses *N* fluctuation and is hence a special case of DDH with extremely strong *N* regulation near *C*_*k*_.

We shall first clarify the first paradox, the “paradox of changing *N*’s”. This paradox is in the ecological time scale whereby *N* is either growing or oscillating around the carrying capacity (**Fig. 2a**). In this time scale, drift may often (but certainly not always) increase in strength as *N* increases. While *N* and *N*_*e*_ are often poorly correlated in WF models(Crow and Kimura 1970; Charlesworth 2009; Lynch, et al. 2016), there is no demonstration how *N* and *N*_*e*_ can be negatively correlated in the absence of *N* regulation. Nevertheless, the WF models do work well in the evolutionary time scale (**Fig. 2b**). For example, by the PSMC model (Li and Durbin 2011), both the European and Chinese populations experience a severe bottleneck 10-60 kyr ago. Presumably, various environmental forces may have reduced *C*_*k*_ drastically. Averaged over the long-time span, *N* should be at or near *C*_*k*_. In short, the N value in the WF models is both N and *C*_*k*_ at the evolutionary time scale. It should also be mentioned that the DDH model is distinct from the concept of “genetic draft” that involves selection and hitchhiking (Gillespie 2000, 2001).

The second paradox of sex-dependent drift is about different *V*(*K*)’s between sexes (generally *V*_*m*_ > *V*_*f*_) but the same *E*(*K*) between them. In the conventional models of sampling, it is not clear what sort of biological sampling scheme could yield *V*(*K*) ≠ *E*(*K*), let alone two separate *V*(*K*)’s with one single *E*(*K*). Mathematically, given separate *K* distributions for males and females, it is unlikely that *E*(*K*) for the whole population could be 1, hence, the population would either explode in size or decline to zero. In short, *N* regulation has to be built into the genetic drift model as the GH model does to avoid this paradox.

The third paradox of genetic drift is manifested in the fixation probability of an advantageous mutation, 2*s*/*V*(*K*). As explained above, the fixation probability is determined by the probability of reaching a low threshold that is independent of *N* itself. Hence, the key parameter of drift in the WF model, *N* (or *N*_*e*_), is missing. This paradox supports the assertion that genetic drift is fundamentally about *V*(*K*) with *N* being a scaling factor. Note the absence of genetic drift at all *N*’s when *V*(*K*) = 0.

The fourth paradox is about the multi-copy gene systems such as viruses and rRNA genes covered in the companion study (Wang, et al. 2024). These systems evolve both within and between hosts. Given the small number of virions transmitted between hosts, drift is strong in both stages as shown by the Haldane model (Ruan, Luo, et al. 2021; Ruan, Wen, et al. 2021; Hou, et al. 2023). Therefore, it does not seem possible to have a single effective population size to account for the genetic drift in two stages with very different biological processes. The inability to deal with multi-copy gene systems may explain the difficulties in accounting for the SARS-CoV-2 evolution (Deng, et al. 2022; Pan, Liu, et al. 2022; Ruan, et al. 2022; Hou, et al. 2023; Ruan, et al. 2023).

As the domain of evolutionary biology expands, many new systems do not have definable populations that fit the criteria of WF populations (such as panmixia in mating or dispersal). Multi-copy gene systems are obvious examples. Others include domestications of animals and plants that are processes of rapid evolution (Diamond 2002; Larson and Fuller 2014; Purugganan 2019; Chen, Yang, et al. 2022; Pan, Zhang, et al. 2022; Wang, et al. 2022). Due to the very large *V*(*K*) in domestication, drift must have played a large role. Somatic cell evolution is another example with “undefinable” genetic drift (Wu, et al. 2016; Chen, et al. 2017; Chen, et al. 2019; Ruan, et al. 2020; Chen, Wu, et al. 2022). The Haldane model is an individual-output model (Chen, et al. 2017) whereby the collection of individuals constitute the “population”.

We understand that further modifications of the WF models may account for some or all of these paradoxes. However, such modifications have to be biologically feasible and, if possible, intuitively straightforward. Such possible elaborations of WF models are beyond the scope of this study. We are only suggesting that the Haldane model can be extensively generalized to be an alternative approach to genetic drift. The GH model attempts to integrate population genetics and ecology and, thus, can be applied to genetic systems far more complex than those studied before. The companion study is one such example.

## Methods

### Cell culture and image analysis

NIH3T3 cells, a fibroblast cell line that was isolated from a mouse NIH/Swiss embryo, were stably transfected with the fluorescent, ubiquitination-based cell cycle indicator (Fucci) (Sakaue-Sawano, et al. 2008) plasmid using Lipofectamine 3000 Transfection Reagent (Invitrogen) following the manufacturer’s specified instructions. The Fucci-labeled cells exhibited distinct fluorescent signals indicative of G1, S, and G2/M phases, represented by red, yellow, and green, respectively. NIH3T3-Fucci cell was derived from single-cell colony and cultured in DMEM supplemented with 10% Calf Bovine Serum and penicillin/streptomycin. All cells were maintained at 37□ with 5% CO_2_. Subsequently, the cells underwent extended time-lapse imaging using high-content fluorescence microscopy (PerkinElmer Operetta CLS) equipped with a 10x objective lens. Images were captured hourly over a 100-hour period, and the analysis was conducted using ImageJ (Fiji) (Schneider, et al. 2012) to count the number of cells (**Supplementary Table 1**).

### Yeast strain construction

Strains were constructed on the genetic background of *Saccharomyces cerevisiae* strain BY4741. A GFP or BFP fluorescent protein expression cassette, under the control of the TDH3 promoter, was inserted into the pseudogene locus YLL017W. Transformations followed a published protocol (Gietz and Schiestl 2007). Transformants were plated on synthetic complete medium without uracil (SC-Ura), and from these, single colonies were selected. Confirmation of replacements was achieved through PCR, and cassette verification was performed using fluorescence microscopy. Subsequently, the constructed strains were cultivated non-selectively in YPD medium (1% Yeast extract, 2% Peptone, 2% D-glucose) at 30 □ on a rotary shaker.

### Estimation of *V*(*K*) and *E*(*K*) in yeast cells

To discern division events, even under high concentration, we conducted co-cultures of the GFP-yeast and BFP-yeast at ratios of 1:1 and 1:25, with the initial cell concentration of 0.1% and 12.5%, respectively. Then the yeast cells were then continuously imaged under high-content fluorescence microscopy (PerkinElmer Operetta CLS) for 10 hours with 1-hour intervals to observe the individual offspring of GFP-yeast. Yeast cells with a distinct offspring number (*K*) within the initial 5 hours (with the high-density group limited to the first 4 hours) were documented (**Supplementary Table 2**). Subsequently, the mean and variance of *K* for each cell were calculated across various time intervals as follows.

According to the law of total variance,

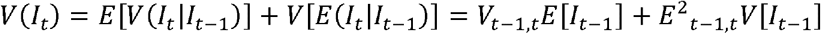

where *I*_*t*_ is the offspring number at time *t* for an initial single cell (i.e., the total number of progeny cells for a single cell after *t* hours), as documented in **Supplementary Table 2**. And *E*_*t*-1, *t*_ represents the average of offspring number after single time interval (1 hour here) for each single cell from time *t*-1 to time *t*, while *V*_*t*-1, *t*_ is the corresponding variance. Utilizing the documented total cell count at time *t* (denoted by *N*_*t*_), we could calculate the average of offspring number for a single yeast cell in a specific time interval, e.g., *E*_*t*-1, *t*_ = *N*_*t*_ / *N*_*t*-1_ and *E*[*I*_*t*-1_] = *N*_*t*_ /*N*_0_. With some rearrangement,

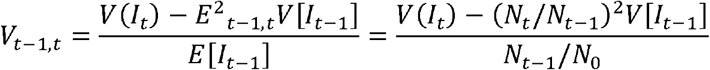

Applying the aforementioned equations, we observed that both *V*_*t*-1, *t*_ (i.e., *V*(*K*) within a one-hour interval) and *V*(*I*_*t*_) (i.e., the *V*(*K*) from 0 to time *t*) for yeast cells under high density (12.5%) are generally greater than the values under low density (**Supplementary Table 3**). This suggests that the variance of offspring number in high-density conditions could be larger than that in a low-density context. It’s noteworthy that the estimated value of *V*_*t*-1, *t*_ could be less than zero (**Supplementary Table 3**), implying a very small variance in the number of offspring. Furthermore, the occurrence of negative values for *V*_*t*-1, *t*_ is more frequent under low density than high density, indicating the higher variance of offspring number in high density as previously suggested.

To show the variance of offspring number overtime, we also calculated the change of offspring number within successive one-hour intervals, denoted as *V*(*I*_*t*_ - *I*_*t*-1_). This value is intricately linked to the variance of offspring overtime. To facilitate the comparison with the total variance from 0h to 4h, we set *V*(*ΔK*) = 4*V*(*I*_*t*_ - *I*_*t*-1_) to account for the four-hour time span.

### Simulation of genetic drift in the Haldane model and the Wright-Fisher (WF) model

In both models, interactions between individuals are implicitly included through the dependency of the average number of offspring on population size, as defined by Eq. (5). This dependency leads to the logistic population growth, reflecting the density-dependent interactions. The initial population size is set to 4, with initial gene frequency (i.e., the frequency of mutant allele) of 0.5. And the carrying capacity is established at 100, 000, with *r* = 1 and *z* = 1. In WF model, the population size at next generation is *N*_*t*+1_ = *N*_*t*_ ×*E*(*K*_*t*_). Note if the calculated value of *N*_*t*+1_ is not an integer, we round it to the nearest whole number. The number of mutant alleles follows a binomial distribution, allowing the simulation of gene frequency overtime in WF model. For Haldane model, the number of offspring number is assumed to follow a beta-binomial distribution. The mean of offspring number *E*(*K*_*t*_) is obtained from Eq. (5). The variance *V*(*K*_*t*_) is obtained from Eq. (6) with parameters *a* = 1/300 and *b* = 1.2. Given the mean and variance of the beta-binomial distribution, we simulated the number of mutant alleles and population size overtime and then traced the gene frequency overtime.

### Fixation probability of a new mutation in Haldane model

Here we obtain the fixation probability of advantageous mutation in Haldane model governed by a branching process. The Haldane model considers a well-mixed population of *N*_*t*_ haploid parents with only two types of alleles (W for wildtype, M for mutant). There is no migration and new mutations in this model. Each individual is assumed to independently reproduce in discrete and non-overlapping generation, *t* = 0, 1, 2, …. Thus, the number of offspring per allele (also an individual in this haploid population) at any generation is represented by a set of identically and individually distributed random variables. In particular, the numbers of offspring of M and W alleles (denoted as *K*_*M*_ and *K*_*W*_ respectively) can be represented by following distributions.

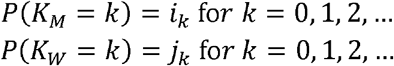

For a particular distribution, we can obtain the average number of offspring as follows.

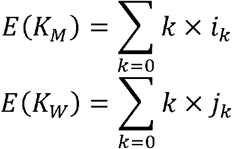

With the selection coefficient of *s* (0 for neutral mutation, positive for advantageous mutation, and negative for deleterious mutation) of M allele, average offspring number of M allele and W allele will follow the relationship.

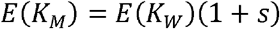

Now, the evolution of the number of M allele (denoted as *M*_*t*_) and the number of W allele (denoted as *W*_*t*_) as time process is a branching process.

To compare Haldane model with WF model (the offspring number follows Poisson distribution with variance and mean equal to 1), we will set *E*(*K*_*W*_) = 1, *E*(*K*_*M*_) = 1 + *s*. And then let the offspring number follow a more general and realistic distribution (including negative binomial distribution (negBin), beta-binomial distribution (betaBin)), which can let the ratio of variance to mean range from 1 to a large number. (For simplicity, we let *V*(*K*_*M*_)/*E*(*K*_*M*_) *= V*(*K*_*W*_)/*E*(*K*_*W*_).) Based on the branching process, we obtained the fixation probability M alleles (see Supplementary Information for more details on the derivation).

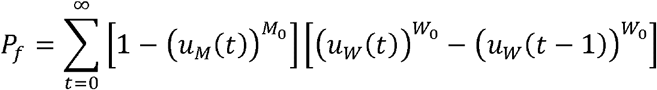

where

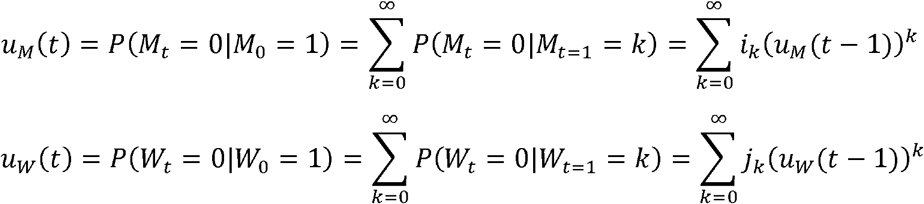

Note *u*_*M*_(*t* = 1) = *P*(*K*_*M*_ = 0) = *i*_0_, *u*_*W*_(*t* = 1) = *P*(*K*_*W*_ = 0) = *j*_0_. And both {*u*_*M*_(*t*), *t* = 0, 1, 2, …} and {*u*_*W*_(*t*), *t* = 0, 1, 2, …} are a bounded monotonic sequence. Thus, although we cannot obtain the direct analytical solution for *P*_*f*_, we could easily obtain their numerical solution by iteration to the case when both *u*_*M*_(*t*) and *u*_*W*_(*t*) converge.

## Acknowledgements

We thank Xionglei He for helpful comments. The work was supported by the National Natural Science Foundation of China (32150006, 32200493, 32293193, 81972691), the Guangzhou Science and Technology Planning Project (2025A04J3499), the National Key Research and Development Projects of the Ministry of Science and Technology of China (2021YFC2301300, 2021YFC0863400).

