## Supplementary Information for "The Generalized Haldane (GH) model tracking population size changes and resolving paradoxes of genetic drift"

### The Generalized Haldane (GH) model of genetic drift resolving the many paradoxes of molecular evolution

Ruan et al.

The file includes:

I. Haldane model and assumptions

II. Density-Dependent Haldane (DDH) model

III. Male-to-female ratio (*α'* = *V_m_*/*V_f_*) of offspring number variance

Supplementary Table 4

Supplementary Figs. 1 to 5

References

Other Supplementary Information for this manuscript includes the following:

Supplementary Table 1 to 3

I. Haldane model and assumptions

The Haldane model considers a well-mixed population of *N_t_* haploid parents with only two types of alleles (M and W allele). However, all the results can be extended to a diploid population, in which mating is entirely at random, with initial population turns to be *N_t_*/2. There is no migration and new mutations in this model. The numbers of M allele and W allele at generation *t* are *M_t_* and *W_t_* respectively. Thus, the frequency of M allele is *x_t_* = *M_t_* /*N_t_*.

Each individual is assumed to independently reproduce in discrete and non-overlapping generation, *t* = 0, 1, 2, …. Note the distribution of offspring number could follow any distribution, e.g., Poisson, negative binomial, and beta-binomial distribution. Thus, the number of offspring per allele (also an individual in this haploid population) at any generation is represented by a set of identically and individually distributed random variables. In particular, the numbers of offspring of M and W alleles (denoted as *K_M_* and *K_W_* respectively) can be represented by following distributions.

$$P\left( K_{M}=k \right)=i_{k} \mathrm{fo}r k=0, 1, 2, \ldots$$

$$P\left( K_{W}=k \right)=j_{k} \mathrm{fo}r k=0, 1, 2, \ldots$$

Then the evolution of the population as time processes, including processes {*M_t_*, *t* = 0, 1, 2, …} and {*W_t_*, *t* = 0, 1, 2, …}, is a branching process. And the fitness of M allele and W allele can be represented by the average of offspring number.

$$E\left( K_{M} \right)=\sum_{k=0} k\times i_{k}$$

$$E\left( K_{W} \right)=\sum_{k=0} k\times j_{k}$$

Let the selection coefficient of M allele be *s* (positive for advantageous mutation, negative for deleterious mutation), offspring number average of M allele and W allele will have the following relationship:

$$E\left( K_{M} \right)=E\left( K_{W} \right)\left( 1+s \right)$$

In Wright-Fisher model, *V*(*K_M_*) = *E*(*K_M_*) and *V*(*K_W_*) = *E*(*K_W_*) based on the assumption that the offspring number follows Poisson distribution (PS). It’s with caveats about its limitations and lack of biological realism. In the Haldane model, the offspring number could be extended to a more realistic distribution, including negative binomial distribution (variance > mean), geometric distribution (variance > mean), beta-binomial distribution (variance ≥ mean), binomial distribution (variance < mean) and so on.

Under the Haldane model governed by branching processes, we will obtain genetic drift, fixation probabilities and fixation times of M allele in the following subsections, which are the key parameters to describe the evolutionary dynamic of a population.

1. Genetic drift in Haldane model

In population genetics, the genetic drift is generally represented by the variance of gene frequency of an allele. According to the Individual Output (IO) model based on branching process theory, Chen et al. obtained the genetic drift for haploid population with some approximation (Chen, et al. 2017).

$$V\left( \Delta x \right)=V\left( x^{'}-x \right)=V(x')=\frac{V(K)}{E^{2}(K)}\frac{x(1-x)}{N}$$

Note the accurate analytical equation has not been derived. And the approximation was obtained from Kimura and Crow (1963) while setting *α* = 1 and *N*_e_ = 1/2*N* (since cell populations are asexual) in their equation (18). In Kimura and Crow’s calculation, they considered the output of each gamete, but they still assumed that the population size is determined externally. And this assumption will lead to the reproduction of each gamete or individual is not independent. Instead, the covariance in progeny number for random pairs from the parental generation is negative (see their Eq. (14)), suggesting the competition, rather that independence, among gamete or individuals. In fact, this is different from the property of “independence among individual” in Haldane model. Here we give an alternative method to obtain the approximate equation based on Haldane model.

The Haldane model considers a well-mixed population of *N* haploid parents with only two types of alleles (wildtype and mutant allele). There is no migration and new mutations in this model. Each individual is assumed to independently reproduce in discrete and non-overlapping generation, with the mean of variance of offspring number (*K*) equal to *E*(*K*) and *V*(*K*). In the absence of selection, the expected frequency of M allele at offspring generation will be the same as that at parental generation.

$$E\left( x^{'} \right)=E\left( \frac{M'}{M'+W'} \right)=x=\frac{M}{M+W}=\frac{M}{N}$$

where *M* and *W* are the number of mutant and wildtype alleles at parental generation respectively. And *M’* and *W’* are the number of mutant and wildtype alleles at offspring generation respectively.

According to Taylor approximations of the ratio of two variable (Kendall 1948; Hubbard and Hubbard 2015), we have the (first-order) approximation of the variance of M allele frequency at offspring generation:

$$V\left( x^{'} \right)=V\left( \frac{M^{'}}{N^{'}} \right)\approx\left( \frac{E\left( M^{'} \right)}{E\left( N^{'} \right)} \right)^{2}\left( \frac{V\left( M^{'} \right)}{E^{2}\left( M^{'} \right)}+\frac{V\left( N^{'} \right)}{E^{2}\left( N^{'} \right)}-\frac{2\mathrm{Cov}(M^{'}, N^{'})}{E\left( M^{'} \right)E\left( N^{'} \right)} \right)$$

Note

$$E\left( M^{'} \right)=ME\left( K \right)$$

$$E\left( N^{'} \right)=NE\left( K \right)$$

$$V\left( M^{'} \right)=MV\left( K \right)$$

$$V\left( N^{'} \right)=NV(K)$$

$$\mathrm{Cov}\left( M^{'}, N^{'} \right)=\mathrm{Cov}\left( M^{'}, M^{'}+W^{'} \right)=\mathrm{Cov}\left( M^{'}, M^{'} \right)+\mathrm{Cov}\left( M^{'}, W^{'} \right)=V\left( M^{'} \right)+0=MV\left( K \right)$$

Thus,

$$V\left( x^{'} \right)\approx\left( x \right)^{2}\left( \frac{MV\left( K \right)}{ME\left( K \right)ME\left( K \right)}+\frac{NV\left( K \right)}{NE\left( K \right)NE\left( K \right)}-\frac{2MV\left( K \right)}{ME\left( K \right)NE\left( K \right)} \right)=\frac{V\left( K \right)}{E^{2}\left( K \right)}\frac{x\left( 1-x \right)}{N} (A1)$$

2. Approximation of fixation probability with the aid of diffusion equations

Based on the diffusion equation (Crow and Kimura 1970), the fixation probability of a mutation with initial frequency of *x* is

$$u\left( x \right)=\frac{1-e^{-4N_{e}sx}}{1-e^{-4N_{e}s}}$$

where *s* is selective coefficient (*s* > 0 and *s* < 0 for selective advantage and dis-advantage, respectively). In the main text, we know that the effective population size in Haldane model is

$$N_{e}=\frac{NE^{2}(K)}{V\left( K \right)}$$

We set *E*(*K*) = 1 for the wildtype allele and *E*(*K*) > 1 for the advantageous allele. Thus, *E*(*K*) for the entire population would be slightly larger than 1. And then we can obtain the approximate fixation probability of new advantageous mutation as follows:

$$u\left( x \right)=\frac{1-e^{-4N_{e}sx}}{1-e^{-4N_{e}s}}\approx\left( 2s \right)2N_{e}x=\left( 2s \right)\frac{2Nx}{V\left( K \right)} (A2)$$

When *s* = 0, *u*(*x*) = *x*. As expected, when all variants have the same fitness, the fixation probability should equal the variant frequency, regardless of the value of *V*(*K*). When *s* > 0, the fixation probability for a new advantageous mutation (*x* = 1/2*N*) can be approximated by

$$u=\frac{2s}{V\left( K \right)} (A3)$$

If *V*(*K*) = *E*(*K*) = 1 as in the WF model, we also confirm the classical result of *u* = 2*s* (Haldane 1932; Crow and Kimura 1970).

3. Fixation probability in Haldane model

At generation *t*, there will be four states for W allele and M allele.

| **Coextinction**  *M_t_* = 0 and *W_t_* = 0 | **M allele extinction**  *M_t_* = 0 and *W_t_* ≥ 1 |
| --- | --- |
| **M allele fixation**  *M_t_* ≥ 1 and *W_t_* = 0 | **Coexistence**  *M_t_* ≥ 1 and *W_t_* ≥ 1 |

Given the offspring number distribution of M allele ($i_{0},i_{1},\ldots, i_{k}$) and W allele ($j_{0},j_{1},\ldots, j_{k}$), we can obtain the probabilities that M, W alleles are extinct by generation *t* while the initial numbers of M allele and W allele are both 1 (i.e., *M*_0_ = 1, *W*_0_ = 1).

$$u_{M}\left( t \right)=P\left( M_{t}=0|M_{0}=1 \right)=\sum_{k=0}^{\infty} P\left( M_{t}=0|M_{t=1}=k \right)=\sum_{k=0}^{\infty} i_{k}\left( u_{M}(t-1) \right)^{k} (A4)$$

$$u_{W}\left( t \right)=P\left( W_{t}=0|W_{0}=1 \right)=\sum_{k=0}^{\infty} P\left( W_{t}=0|W_{t=1}=k \right)=\sum_{k=0}^{\infty} j_{k}\left( u_{W}(t-1) \right)^{k} (A5)$$

where $u_{M}\left( t=1 \right)=i_{0}=P(K_{M}=0)$,$u_{W}\left( t=1 \right)=j_{0}=P(K_{W}=0)$. Note $u_{M}\left( t \right)\leq u_{M}\left( t+1 \right)$ and $u_{M}\left( t+1 \right)\leq1$ – i.e., {$u_{M}\left( t \right)$} is a bounded monotonic sequence. Likewise, {$u_{W}\left( t \right)$} is also a bounded monotonic sequence. As generation t proceeds, both $u_{M}\left( t \right)$ and $u_{W}\left( t \right)$ will converge to a particular value, which is called the probability of ultimate extinction. Thus, we can obtain the ultimate probability that M allele or W allele is extinct conditioned on the initial number of M allele or W allele is 1.

$$u_{M}=\lim_{t\to\infty} u_{M}\left( t \right)$$

$$u_{W}=\lim_{t\to\infty} u_{W}\left( t \right)$$

Based on the definition of the Haldane model, we know that the offspring numbers of M or W (denoted as *K_M_*, *K_W_* respectively) follows a particular distribution. And their probability generating function (PGF) could be defined as following.

$$G_{M}\left( s \right)=\sum_{k=0}^{\infty} i_{k}s^{k}$$

$$G_{W}\left( s \right)=\sum_{k=0}^{\infty} j_{k}s^{k}$$

According to the properties of probability generating function (Grimmett and Stirzaker 2009), we can also obtain the PGF of *M_t_* and *W_t_* by the *t*-fold iterate as following.

$$G_{M_{t}}\left( s \right)=G_{M}\left( G_{M_{t-1}}\left( s \right) \right)=G_{M}\left( G_{M}\left( G_{M_{t-2}}\left( s \right) \right) \right)=G_{M}\left( G_{M}\left( \ldots\left( G_{M}\left( s \right) \right)\ldots\right) \right)$$

$$G_{W_{t}}\left( s \right)=G_{W}\left( G_{W_{t-1}}\left( s \right) \right)=G_{W}\left( G_{W}\left( G_{W_{t-2}}\left( s \right) \right) \right)=G_{W}\left( G_{W}\left( \ldots\left( G_{W}\left( s \right) \right)\ldots\right) \right)$$

And then

$$u_{M}\left( t \right)=P\left( M_{t}=0|M_{0}=1 \right)=G_{M_{t}}\left( s=0 \right)=G_{M}\left( G_{M}\left( \ldots\left( G_{M}\left( s=0 \right) \right)\ldots\right) \right)$$

$$u_{W}\left( t \right)=P\left( W_{t}=0|W_{0}=1 \right)=G_{W_{t}}\left( s=0 \right)=G_{W}\left( G_{W}\left( \ldots\left( G_{W}\left( s=0 \right) \right)\ldots\right) \right)$$

Each allele gives birth to offspring independently. Thus,

$$P\left( M_{t}=0|M_{0} \right)=\left( u_{M}\left( t \right) \right)^{M_{0}}$$

$$P\left( W_{t}=0|W_{0} \right)=\left( u_{W}\left( t \right) \right)^{W_{0}}$$

In population genetics, the fixation of M allele means that the population consist entirely of the M allele, with no W alleles remaining. We define the fixation probability of M allele by generation *t* as follows:

$$P_{10}\left( t \right)=P(M_{t}\geq1, W_{t}=0|M_{0},W_{0})$$

Given that M and W allele reproduce independently, this can be factored as:

$$P_{10}\left( t \right)=P\left( M_{t}\geq1|M_{0} \right)P\left( W\left( t \right)=0|W_{0} \right) =\left[ 1-\left( u_{M}\left( t \right) \right)^{M_{0}} \right]\left[ \left( u_{W}\left( t \right) \right)^{W_{0}} \right] (A6)$$

As *t* approaches infinity, the ultimate fixation probability of M allele can be derived as follows:

$$P_{10}=\lim_{t\to\infty} P_{10}\left( t \right)=\lim_{t\to\infty} P\left( M_{t}\geq1 \right)P\left( W_{t}=0 \right) =\left( 1-{u_{M}}^{M_{0}} \right){u_{W}}^{W_{0}} (A7)$$

Note when M allele is fixed at generation *t*, there will be two cases in subsequent generations: M allele keep fixed or coextinction of two alleles. Thus, the fixation state of M allele at generation *t* includes two independent cases: (i) *M_t_* ≥ 1 and *W_t_* = 0, *W_t_*_-1_ = 0, i.e., M allele is fixed at generation *t* – 1 and it keeps fixed at generation *t*; (ii) *M_t_* ≥ 1 and *W_t_* = 0, *W_t_*_-1_ ≥ 1, i.e., M allele is fixed for the first time at generation *t* (M allele and W allele coexist at generation *t* – 1, and then only the W allele turn extinct at generation *t*). And the probability that M allele is fixed for the first time at generation *t* is

$$F_{10}\left( t \right)=P\left( M_{t}\geq1 \right)P\left( W_{t}=0,W_{t-1}\geq1 \right)=P\left( M_{t}\geq1 \right)\left[ P\left( W_{t}=0 \right)-P\left( W_{t-1}=0 \right) \right] =\left[ 1-\left( u_{M}\left( t \right) \right)^{M_{0}} \right]\left[ \left( u_{W}\left( t \right) \right)^{W_{0}}-\left( u_{W}\left( t-1 \right) \right)^{W_{0}} \right] (A8)$$

The cumulative probability that M allele is fixed for the first time over time is

$$F_{10}=\sum_{t=0}^{\infty} F_{10}\left( t \right)=\sum_{t=0}^{\infty} \left[ 1-\left( u_{M}\left( t \right) \right)^{M_{0}} \right]\left[ \left( u_{W}\left( t \right) \right)^{W_{0}}-\left( u_{W}\left( t-1 \right) \right)^{W_{0}} \right] (A9)$$

Similarly, the loss probability of M allele at generation *t* (i.e., *M_t_* = 0 and *W_t_* ≥ 1) and the ultimate loss probability of M allele are

$$P_{01}\left( t \right)=\left[ \left( u_{M}\left( t \right) \right)^{M_{0}} \right]\left[ 1-\left( u_{W}\left( t \right) \right)^{W_{0}} \right] (A10)$$

$$P_{01}=\lim_{t\to\infty} P_{01}\left( t \right)=\lim_{t\to\infty} P\left( M_{t}=0 \right)P\left( W_{t}\geq1 \right) ={u_{M}}^{M_{0}} \left( 1-{u_{W}}^{W_{0}} \right) (A11)$$

The probability that M allele is lost for the first time at generation *t* (i.e., *M_t_* = 0| *M_t_*_-1_ ≥ 1 and *W_t_* ≥ 1) and the cumulative probability that M allele is lost for the first time over time are

$$F_{01}\left( t \right)=\left[ \left( u_{M}\left( t \right) \right)^{M_{0}}-\left( u_{M}\left( t-1 \right) \right)^{M_{0}} \right]\left[ 1-\left( u_{W}\left( t \right) \right)^{W_{0}} \right] (A12)$$

$$F_{01}=\sum_{t=0}^{\infty} F_{01}\left( t \right)=\sum_{t=0}^{\infty} \left[ \left( u_{M}\left( t \right) \right)^{M_{0}}-\left( u_{M}\left( t-1 \right) \right)^{M_{0}} \right]\left[ 1-\left( u_{W}\left( t \right) \right)^{W_{0}} \right] (A13)$$

Note that there is a certain probability that the population will be extinct ultimately, i.e., coextinction of both M allele and W allele,

$$P_{00}=\lim_{t\to\infty} P_{00}\left( t \right)=\lim_{t\to\infty} P\left( M_{t}=0 \right)P\left( W_{t}=0 \right) ={u_{M}}^{M_{0}}{u_{W}}^{W_{0}}$$

Thus, even though one of the two alleles is fixed at generation *t*, the number of this allele could still turn to be zero in subsequence generations. That’s to say, an allele cannot always remain fixed because of the coextinction of both alleles, which leads to *F*_10_ ≥ *P*_10_, *F*_01_ ≥ *P*_01_. However, if additionally assume the population will not be extinct (this could be achieved by simply let the population size keeps around a certain number with the introduction of density-dependent branching processes, i.e., the density dependent Haldane model in the next section), then *F*_10_ = *P*_10_, *F*_01_ = *P*_01_. Now the fixation probability and loss probability of M allele are

$$P_{f}=F_{10}=\sum_{t=0}^{\infty} F_{10}\left( t \right)=\sum_{t=0}^{\infty} \left[ 1-\left( u_{M}\left( t \right) \right)^{M_{0}} \right]\left[ \left( u_{W}\left( t \right) \right)^{W_{0}}-\left( u_{W}\left( t-1 \right) \right)^{W_{0}} \right] (A14)$$

$$P_{e}=F_{01}=\sum_{t=0}^{\infty} F_{01}\left( t \right)=\sum_{t=0}^{\infty} \left[ \left( u_{M}\left( t \right) \right)^{M_{0}}-\left( u_{M}\left( t-1 \right) \right)^{M_{0}} \right]\left[ 1-\left( u_{W}\left( t \right) \right)^{W_{0}} \right] (A15)$$

In this case, we could obtain the average fixation time, the average loss time of M allele.

$$T_{1}=\frac{1}{F_{10}}\sum_{t=1}^{\infty} t\times F_{10}\left( t \right) (A16)$$

$$T_{0}=\frac{1}{F_{01}}\sum_{t=1}^{\infty} t\times F_{01}\left( t \right) (A17)$$

As mentioned before, both $u_{M}\left( t \right)$ and $u_{W}\left( t \right)$ will converge to a particular value as time *t* increases, which will lead that *F*_01_(*t*) and *F*_10_(*t*) will converge to 0 while *t* is large enough. And then *P_f_*, *P_e_*, *T*_1_, and *T*_0_ will converge to a particular value. Thus, although we cannot obtain the direct analytical solution for *P_f_*, *P_e_*, *T*_1_, and *T*_0_, we could easily obtain their numerical solution by iteration to the case when both *F*_01_(*t*) and *F*_10_(*t*) converge to 0.

4. Confirmation the accuracy of the fixation probability using simulation

We have confirmed the accuracy of both the approximate fixation probability inferred from Eq. (A3) and the numerical solution from Eq. (A14), using the simulation based on the branching process (**Supplementary Fig. 4**). The numerical solution from Eq. (A14) is the same as the simulation result. Moreover, the approximation of Eq. (A3) is sufficiently accurate as long as *N*≥50 which is, for all practical purposes, quite adequate. When *N* is as small as 10, the theoretical result is biased. Indeed, at such a low *N* value, the population is prone to extinction. Similar views have been expressed by Burden and Simon (2016) who, however, did not model selection.

The result of *N* = 10 agrees with the common criticisms against the branching process in Haldane model, whereby the population size is intrinsically unstable. In the branching process, *E*(*K*) = 1 would mean constant population size averaged over all populations. However, each individual population would either go extinct or approach infinite size in the branching process. While this instability in *N* has been considered an undesirable property for modeling genetic drift (Burden and Simon 2016). We shall rectify this intrinsic property of instability by incorporating the regulation of *N* into the model. Such a model will be presented in the next section.

II. Density-Dependent Haldane (DDH) model

The impactful limitation of the WF model is that it imposes *N* externally, rather than generating *N* from within the model. In contrast, the branching process can incorporate *N* regulation and would be a more general model for genetic drift than the WF model. Since *N* would approach zero or infinity in the branching process without regulating *N*, correcting this deficiency is both necessary and beneficial. This modification is referred to as the density-dependent Haldane model (DDH).

1. The structure of the density-dependent Haldane (DDH) model

Here we consider a haploid population with two neutral alleles (W for wildtype, M for mutant). The population size at time *t* is *N_t_* among which *M_t_* are the mutants. The frequency of the M allele at generation *t* is *x_t_* = *M_t_*/*N_t_*. The growth of populations is often density-dependent due to competition for limited resources. The density-dependent branching process is defined by the following recursive equation.

$$E\left( K_{t} \right)=\frac{E\left( N_{t+1} \right)}{N_{t}}=\frac{r+1}{1+r\left( \frac{N_{t}}{C_{k}} \right)^{z}} (A18)$$

Here *r* is the intrinsic growth rate, i.e., the proportional increase of the population *N_t_* in one unit of time at the early, unimpeded stage (*N_t+_*_1_ - *N_t_* = *rN_t_*). When there is no competition for the resources (i.e., *N_t_* is close to 0), *E*(*K_t_*) = *r* + 1. When the population size reaches carrying capacity, *E*(*K_t_*) = 1. The parameter *z* determines the curvature of *N_t_* over time. And the population dynamics could be modelled as follows.

$$N_{t+1}=\sum_{i=1}^{N_{t}} K_{t}^{\left( i \right)}\left( C_{k},N_{t} \right) (A19)$$

where *K_t_*^(^*^i^*^)^(*C_k_*, *N_t_*) is a generical random variable that represents the number of offspring of the *i*-th individual (or allele) with the carrying capacity *C_k_* and the population size *N_t_* at generation *t*. Without selection effect, *K_t_*^(^*^i^*^)^(*C_k_*, *N_t_*) is identically distributed random variable for all *i*, simply denoted as *K_t_*. And the mean and variance of offspring numbers of each individual at generation *t* are

$$E\left( K_{t} \right)=\sum_{k=0} k\times P(K_{t}=k) (A20)$$

$$V\left( K_{t} \right)=\sum_{k=0} \left[ E\left( K_{t} \right)-k \right]^{2}\times P(K_{t}=k) (A21)$$

Then the evolution of the population as time processes, {*N_t_*, *t* = 0, 1, 2, …} and {*M_t_*, *t* = 0, 1, 2, …}, is a branching process.

2. Negative binomial distribution of K - The biological reasoning

Based on the overdispersion of offspring number (Table 1), we assume the offspring number follows negative binomial distribution (*V*(*K_t_*) is generally greater than *E*(*K_t_*)) with shape parameters of *n_t_* and *p_t_*. For an individual at generation *t*, the individual reproduces an offspring number at a regular interval. During each reproduction, it would suffer an injury with a probability of *p_t_*. And by *n_t_*-th injury, the individual will die or be infertile. There is an alternative interpretation of the shape parameters: *n_t_* represents the times an individual could reproduce per generation, and 1/*p_t_* represents the offspring number of each reproduction. We use the first interpretation unless otherwise specified. The mean and variance of offspring number will be

$$E\left( K_{t} \right)=\frac{n_{t}}{p_{t}} (A22)$$

$$V\left( K_{t} \right)=\frac{n_{t}(1-p_{t})}{{[p_{t}]}^{2}} (A23)$$

Note because of the density-dependent branching growth resulting from limited resources (defined as Eq. (A18)), both *n_t_* and *p_t_* will depend on population size *N_t_*. Thus,

$$E\left( K_{t} \right)=\frac{E\left( N_{t+1} \right)}{N_{t}}=\frac{n_{t}}{p_{t}}=\frac{r+1}{1+r\left( \frac{N_{t}}{C_{k}} \right)^{z}} (A24)$$

Note the ratio of variance to mean of offspring number may or may not change with the growth of population. Here we simply assume its ratio changes as follows.

$$\frac{V\left( K_{t} \right)}{E\left( K_{t} \right)}=\frac{1-p_{t}}{p_{t}}=a\left( N_{t} \right)^{b} (A25)$$

In this case, the ratio of *V*(*K_t_*)/*E*(*K_t_*) could be constant, decrease or increase with the increase of population size.

With some transformation,

$$p_{t}=\frac{1}{1+a\left( N_{t} \right)^{b}} (A26)$$

$$n_{t}=E\left( K_{t} \right)p_{t}=\frac{r+1}{1+r\left( \frac{N_{t}}{C_{k}} \right)^{z}}\frac{1}{1+a\left( N_{t} \right)^{b}} (A27)$$

3. Genetic drift under the DDH model - the formulation

According to Eq. (A1), we can obtain the genetic drift in general Haldane model after a single generation. Specifically, the variance of M allele frequency is given by:

$$V\left( x_{t+1} \right)=V\left( \frac{M_{t+1}}{N_{t+1}} \right)=\frac{x_{t}\left( 1-x_{t} \right)}{N_{t}}\frac{V\left( K_{t} \right)}{E^{2}(K_{t})} (A28)$$

Note we would like to clarify that Eq. (A1) and Eq. (A28) are essentially the same, with the only difference being the subscript 𝑡, which indicates the time dependence in the dynamic process. And the strength of genetic drift (i.e., normalized by (1- *x_t_*)*x_t_*) is

$$G_{t}=\frac{1}{N_{t}}\frac{V\left( K_{t} \right)}{E^{2}\left( K_{t} \right)}=\frac{1}{N_{t}}\frac{1-p_{t}}{n_{t}}=\frac{1}{N_{t}}\frac{a\left( N_{t} \right)^{b}\left[ 1+r\left( \frac{N_{t}}{C_{k}} \right)^{z} \right]}{r+1} (A29)$$

Where *z* > 0 (*z* = 0 will let *E*(*K_t_*) always equal to 1), *a* ≥ 1. According to the new definition of genetic drift under DDH model, we could obtain the effective population size as following equation.

$$N_{e}=\frac{1}{G_{t}}=\frac{N_{t}(r+1)}{a\left( N_{t} \right)^{b}\left[ 1+r\left( \frac{N_{t}}{C_{k}} \right)^{z} \right]} (A30)$$

The partial derivative of Eq. (A29) with respective to population size *N_t_* is

$${G_{t}}^{'}=\frac{a}{r+1}\left[ \left( b-1 \right)\left( N_{t} \right)^{b-2}+\frac{r(z+b-1)}{\left( C_{k} \right)^{z}}\left( N_{t} \right)^{z+b-2} \right] (A31)$$

When *b* ≥ 1, *G_t_’* will always be greater than 0 and the strength of genetic drift will increase as the growth of the population size. Likewise, when *b* < 1 and z + *b* - 1 ≤ 0, the strength of genetic drift will keep decreasing as the growth of population. However, when *b* < 1 and z + *b* - 1 > 0, the strength of genetic drift will be smallest when population size fits following equation.

$$N_{t}=\left[ \frac{1-b}{r(z+b-1)} \right]^{\frac{1}{z}}C_{k} (A32)$$

As shown, the strength of genetic drift will increase when the population size is either less or greater than *N_t_* of Eq. (A32).

4. DDH modeling of genetic drift as a function of N

We now evaluate the strength of genetic drift in populations with *N* near the carrying capacity *C_k_*, based on the mathematics presented above. In the WF model, *N* is constant at (or near) *C_k_* with *E*(*K*) = 1 = *V*(*K*). Hence, genetic drift in the WF model is solely driven by sampling stochasticity. In contrast, the DDH model incorporates both *E*(*K*) and *V*(*K*) as functions of *N*, making them density dependent.

In traditional Wright-Fisher model, the offspring number *K_t_* is assumed to follow Poisson distribution (PS) with *V*(*K_t_*) always being equal to *E*(*K_t_*). Based on this assumption, the strength of genetic drift of Wright-Fisher model will be 1/*N_t_*. Thus, neutral genetic drift should decrease with increasing population size (*N*). However, the genetic drift redefined by the density-dependent Haldane model (DDH) could result in very different outcomes.

Even though we keep *V*(*K_t_*)/*E*(*K_t_*) ratio equal to 1 as WF model, the strength of genetic drift can indeed increase, decrease or stay nearly constant as population size *N_t_* changes (**Supplementary Fig. 1**). That’s to say, the effective population size could indeed decrease, increase or stay nearly constant as the growth of population size. Moreover, we can see the same pattern regardless of whether the *V*(*K_t_*)/*E*(*K_t_*) ratio increases (**Supplementary Fig. 2**) or decreases (**Supplementary Fig. 3**) as the growth of the population. These results shows that the genetic drift cannot be simply determined by population size. Instead, it depends on the ecology that governs *E*(*K_t_*), *V*(*K_t_*) and *N_t_*.

In summary, genetic drift effect can indeed decrease, increase or stay nearly constant as the population size increases, depending on the strength of *N* regulation near *C_k_*. These results show that the strength of genetic drift depends on the ecology that governs *E*(*K*), *V*(*K*) and *N*. This ecology-based model of genetic drift may more realistically reflect the influences of changing population size on the neutral trajectory of evolution.

III. Male-to-female ratio (*α'* = *V_m_*/*V_f_*) of offspring number variance

In a sexual species, each male or female may produce *K* progeny. The mean of *E*(*K*) should be the same between sexes but the variances, *V_m_* and *V_f_*, could be very different. While the *K* distribution, hence *V_f_*, is observable in females, *V_m_* is rarely known. The variance, *V_m_* and *V_f_*, has direct effect on the genetic drift. Thus, their genetic diversities can inform about the ratio of *α'* = *V_m_*/*V_f_*. Here, we surveyed these diversities in humans and other apes. The estimated *α'* based on mean square error in Great Apes is often ≥ 5. When *α'* exceeds 10, the resolution significantly decreases due to its rapid approach towards infinity. In Great Apes, the only exception is bonobo which, curiously, has *α'* < 1 (~ 0.57). Interpretations of chromosome-level diversities involving positive and negative selection are discussed. The theory is conceptually general as *V*(*K*) is the basis of genetic drift in the Haldane model of the branching process. Furthermore, the estimation of *α'* should be broadly useful in behavioral studies.

1. Theory for the estimation of α'

According to the coalescent theory (Kingman 1982b, a; Hudson 1983; Tajima 1983), we could obtain the average number of nucleotide differences between pairwise sequences.

$$\pi=\theta=4N_{e}\mu=4\frac{N}{V\left( K \right)}\mu$$

Applying it to autosome, X, and Y chromosomes,

$$\theta_{A}=4\frac{N_{A}}{V_{A}}\mu_{A},\theta_{Y}=4\frac{N_{Y}}{V_{Y}}\mu_{Y},\theta_{X}=4\frac{N_{X}}{V_{X}}\mu_{X}$$

With some transformation (note *N_A_*: *N_X_*: *N_Y_* = 4:3:1 in human and other apes),

$$\frac{\theta_{Y}}{\theta_{A}}=\frac{N_{Y}}{N_{A}}\frac{V_{A}}{V_{Y}}\frac{\mu_{Y}}{\mu_{A}}=\frac{1}{4}\frac{V_{A}}{V_{Y}}\frac{\mu_{Y}}{\mu_{A}}$$

$$\frac{\theta_{X}}{\theta_{A}}=\frac{N_{X}}{N_{A}}\frac{V_{A}}{V_{X}}\frac{\mu_{X}}{\mu_{A}}=\frac{3}{4}\frac{V_{A}}{V_{X}}\frac{\mu_{X}}{\mu_{A}}$$

$$\frac{\theta_{Y}}{\theta_{X}}=\frac{N_{Y}}{N_{X}}\frac{V_{X}}{V_{Y}}\frac{\mu_{Y}}{\mu_{X}}=\frac{1}{3}\frac{V_{X}}{V_{Y}}\frac{\mu_{Y}}{\mu_{X}}$$

Assuming the mutation rate in females per generation is *μ_f_*, and the relative mutation rate of males to females is *α*, i.e., *μ_m_* = *αμ_f_*. And then

$$\mu_{A}=\frac{1}{2}\mu_{m}+\frac{1}{2}\mu_{f}=\frac{1}{2}\left( \alpha+1 \right)\mu_{f}$$

$$\mu_{Y}=\mu_{m}=\alpha\mu_{f}$$

$$\mu_{X}=\frac{1}{3}\mu_{m}+\frac{2}{3}\mu_{f}=\frac{1}{3}\left( \alpha+2 \right)\mu_{f}$$

$$\mu_{Y}:\mu_{X}:\mu_{A}=\alpha:\frac{\alpha+2}{3}:\frac{\alpha+1}{2}=\frac{2\alpha}{\alpha+1}:\frac{2\left( \alpha+2 \right)}{3\left( \alpha+1 \right)}:1$$

Note the direct mutation rates (i.e., *μ_m_*, *μ_f_*, *μ_A_*, *μ_Y_*, *μ_A_*) are not easy to obtain unless with the available parent-offspring sequencing data. Instead, we could obtain their pairwise ratios based on divergence over a long evolutionary time as Makova and Li (2002) done. (Divergence is equal to twice the mutation rate times time.)

Assuming variance of offspring number of females per generation is *V_f_*, and the relative variance of males to females is *α'*, i.e., *V_m_* = *α'V_f_*. Likewise,

$$V_{A}=\frac{1}{2}V_{m}+\frac{1}{2}V_{f}=\frac{1}{2}\left( \alpha^{'}+1 \right)V_{f}$$

$$V_{Y}=V_{m}=\alpha^{'}V_{f}$$

$$V_{X}=\frac{1}{3}V_{m}+\frac{2}{3}V_{f}=\frac{1}{3}\left( \alpha'+2 \right)V_{f}$$

With some substitutions,

$$\frac{\theta_{Y}}{\theta_{A}}=\frac{N_{Y}}{N_{A}}\frac{V_{A}}{V_{Y}}\frac{\mu_{Y}}{\mu_{A}}=\frac{1}{4}\frac{V_{A}}{V_{Y}}\frac{\mu_{Y}}{\mu_{A}}=\frac{1}{4}\frac{\alpha^{'}+1}{2\alpha^{'}}\frac{2\alpha}{\alpha+1}=\frac{1}{4}\frac{\alpha^{'}+1}{\alpha^{'}}\frac{\alpha}{\alpha+1}$$

$$\frac{\theta_{X}}{\theta_{A}}=\frac{N_{X}}{N_{A}}\frac{V_{A}}{V_{X}}\frac{\mu_{X}}{\mu_{A}}=\frac{3}{4}\frac{V_{A}}{V_{X}}\frac{\mu_{X}}{\mu_{A}}=\frac{3}{4}\frac{3(\alpha^{'}+1)}{2(\alpha^{'}+2)}\frac{2(\alpha+2)}{3(\alpha+1)}=\frac{3}{4}\frac{(\alpha^{'}+1)}{(\alpha^{'}+2)}\frac{(\alpha+2)}{(\alpha+1)}$$

$$\frac{\theta_{Y}}{\theta_{X}}=\frac{N_{Y}}{N_{X}}\frac{V_{X}}{V_{Y}}\frac{\mu_{Y}}{\mu_{X}}=\frac{1}{3}\frac{V_{X}}{V_{Y}}\frac{\mu_{Y}}{\mu_{X}}=\frac{1}{3}\frac{\alpha^{'}+2}{3\alpha^{'}}\frac{3\alpha}{\alpha+2}=\frac{1}{3}\frac{\alpha^{'}+2}{\alpha^{'}}\frac{\alpha}{\alpha+2}$$

With further transformations,

$$\alpha^{'}=\frac{1}{8\frac{\theta_{Y}}{\theta_{A}}\frac{\mu_{A}}{\mu_{Y}}-1}=\frac{1}{4\frac{\theta_{Y}}{\theta_{A}}\frac{\alpha+1}{\alpha}-1} (A34.1)$$

$$\alpha^{'}=\frac{1}{1-\frac{8}{9}\frac{\theta_{X}}{\theta_{A}}\frac{\mu_{A}}{\mu_{X}}}-2=\frac{1}{1-\frac{4}{3}\frac{\theta_{X}}{\theta_{A}}\frac{\alpha+1}{\alpha+2}}-2 (A34.2)$$

$$\alpha^{'}=\frac{2}{9\frac{\theta_{Y}}{\theta_{X}}\frac{\mu_{X}}{\mu_{Y}}-1}=\frac{2}{3\frac{\theta_{Y}}{\theta_{X}}\frac{\alpha+2}{\alpha}-1} (A34.3)$$

Furthermore, let *R_YX_* = *θ_Y_* /*θ_X_*, *R_YA_* = *θ_Y_* /*θ_A_*, *R_XA_* = *θ_X_* /*θ_A_*, *y* = *μ_Y_* /*μ_A_*, and *x* = *μ_X_* /*μ_A_*, we could obtain more simple equations as follows.

$$\alpha^{'}=\frac{1}{8R_{YA}/y-1}= \frac{y}{8R_{YA}-y} (A35.1)$$

$$\alpha^{'}=\frac{1}{1-\frac{8}{9}R_{XA}/x}-2=\frac{16R_{XA}-9x}{9x-8R_{XA}} (A35.2)$$

$$\alpha^{'}=\frac{2}{9R_{YX}\frac{x}{y}-1}=\frac{2y}{9xR_{YX}-y} (A35.3)$$

According to the aforementioned equations, we know that the *α'* value will be smaller as the increase of *R_YA_* (= *θ_Y_* /*θ_A_*) or *R_YX_* (=*θ_Y_* /*θ_X_*), but it will be larger with the increase of *R_XA_* (= *θ_X_* /*θ_A_*).

The male-to-female ratio of mutation rate *α* is 5.25 as estimated by Makova and Li (2002). Setting *α* = 5.25, then *μ_Y_*:*μ_X_*:*μ_A_* = 1.68:0.77:1. That’s to say, *y* = *μ_Y_* /*μ_A_* = 1.68 and *x* = *μ_X_* /*μ_A_* = 0.77. Combining the diversity data (Gao, et al. 2023) of A, X, and Y chromosome in human (*θ_A_* = 6.18×10^-4^, *θ_X_* = 3.92×10^-4^, *θ_Y_* = 1.11×10^-4^), we could obtain the male-to-female ratio of offspring number variance.

$$\alpha^{'}= \frac{y}{8R_{YA}-y}=-6.91$$

$$\alpha^{'}=\frac{16R_{XA}-9x}{9x-8R_{XA}}=1.69$$

$$\alpha^{'}=\frac{2y}{9xR_{YX}-y} =11.55$$

The negative value of *α'* make no sense in biology meaning. And this will be solved by introducing the mean squared error (MSE) in next section.

2. Statistical approximation for the estimation of α'

Following the calculations outlined in Eqs. (A35.1-A35.3) from the previous section, we have derived three methods for estimating the male-to-female ratio of variance of offspring number (*α'* = *V_m_*/*V_f_*) based on the pairwise ratios of the diversities of A, X, and Y chromosomes (*R_YA_, R_XA_,* and *R_YX_*). Specifically, the values of *α'* calculated by these three methods are referred to as *α'_YA_*, *α'_XA_*, and *α'_YX_*, corresponding to the utilization of *R_YA_, R_XA_,* and *R_YX_* respectively. And in some cases, the estimated *α'* could be negative value based on the observed ratios, which make no sense in biological meaning. The unexpected cases result from the two possible scenarios in neutral evolution (although selection may could also lead to these cases): i) the observed diversity ratios of *R_YA_* (= *θ_Y_* /*θ_A_*) or *R_YX_* (=*θ_Y_* /*θ_X_*) are only slightly lower than expected values (denoted as $\bar{R_{YA}}$ and $\bar{R_{YX}}$ respectively). ii) the observed diversity ratio of *R_XA_* (= *θ_X_* /*θ_A_*) is slightly higher than expected value (denoted as $\bar{R_{XA}}$). In fact, the biological meaning of negative *α'_YA_* or *α'_YX_* is that the male-to-female ratio of offspring number variance (*α'* = *V_m_*/*V_f_*) is approximate to infinity (i.e., *V_m_* ≫ *V_f_*). While the negative value for *α'_XA_* has two possible meaning due to non-monotonic function of *x* and *R_XA_*: i) One is that *α'* is infinity (i.e., *V_m_* ≫ *V_f_*) when *α'_XA_* ≤ -2; ii) the other is that *α'* is approximate to zero (i.e., *V_m_*/*V_f_* ~ 0) when -1 ≤ *α'_XA_* < 0 (see Supplementary Fig. 5). Supplementary Fig. 5 shows that the parameter region for *α'* falling between 10 and infinity is very small.

To address the possible discrepancy among the three estimates (*α'_YA_*, *α'_XA_*, and *α'_YX_*) and determine the optimal estimate for *α'*, we employ the mean squared error (MSE) for every conceivable value of *α'*. And the value associated with the minimum MSE is considered as the most accurate estimate for *α'*. Specifically, we generated a list of possible for *α'* values ranging from 0 to 1000 in increments of 0.01 (0, 0.01, 0.02, …, 1000). For each *α'* in the list, we calculated the three expected diversity ratios, $\bar{R_{YA}}$, $\bar{R_{XA}}$, and $\bar{R_{YX}}$ using Eqs. (A35.1-A35.3) with the mutation rate ratios for Y/A (*y* = 1.68) and X/A (*x* = 0.77) from Makova and Li (2002). Given the observed diversity ratios (*R_YA_, R_XA_,* and *R_YX_*) and the expected diversity ratios ($\bar{R_{YA}}$, $\bar{R_{XA}}$, and $\bar{R_{YX}}$), we could calculate the minimum mean squared error (MSE) for each *α'* value as follows:

$$\mathrm{MSE}=\frac{1}{3}\left[ {(R_{YA}- \bar{R_{YA}})}^{2}+{(R_{XA}-\bar{R_{XA}})}^{2} +{(R_{YX}-\bar{R_{YX}})}^{2} \right]$$

The possible value of *α'* yielding the lowest MSE is considered as the best estimate for *α'*, denoted as *α'* (MSE). We found that it’s almost always true that *α'* > 5 in primates. Among primates surveyed, bonobo is the only exception with *α'* < 1 (Table 2). While chimpanzees and bonobos are each's closest relatives, their sexual behaviors are very divergent (de Waal 1995; De Waal and Lanting 2023).

3. Estimation of the diversity of Y, X, and A chromosomes

There are many measures for the within-species genetic diversity, *θ = 4N_e_μ*. Under strict neutrality and demographic equilibrium, these measures should all converge. Here we try to extract the polymorphism data in estimating the diversity for Y, X, and A chromosomes (denoted as *θ_Y_*, *θ_X_*, and *θ_A_*) by minimizing the impact of selection.

In the neutral equilibrium, the infinite site model dictates the frequency spectrum to be *ξ_i_* = *θ*/*i*, where *ξ_i_* is the number of sites with the variant occurring *i* times in *n* samples. Since every frequency bin is a measure of *θ*, different measures put different weights on the *i*-th bin (Fay and Wu 2000; Fu 2022). While *π*, the mean pairwise differences between sequences, is most commonly used in the literature, we use several statistics to minimize the possible influences of selection and demography (Wang et al. 2024). In addition, Watterson estimator (*θ_w_*), based on the number of segregating sites *S_n_*, is also a widely-used measure of the "population mutation rate" (i.e., *θ = 4N_e_μ*) (Watterson 1975).

$$\theta_{w}=S_{n}/a_{n}=S_{n}/\left[ 1+\frac{1}{2}+\frac{1}{3}+\ldots+\frac{1}{n-1} \right]$$

In this study, we used to the Watterson estimator *θ_w_* to estimate the genetic diversities (*θ_Y_*, *θ_X_*, and *θ_A_*) for Y, X, and autosomes, which is similar to the value estimated by nucleotide diversity *π* (see Supplementary Table 4).

Supplementary Table 4

**Supplementary Table 4. Summary of genetic diversity of sex chromosomes and autosomes in Great Apes.**

|  | **Y Chromosome ^a^** | | | |  | **X Chromosome ^a^** | | | |  | **Autosomes ^b^** | | | |
| --- | --- | --- | --- | --- | --- | --- | --- | --- | --- | --- | --- | --- | --- | --- |
|  | ***n*** | ***S*** | **Total bp** | ***θ_w_* (×10^-3^)** | ***π* (×10^-3^)** | ***n*** | ***S*** | **Total bp** | ***θ_w_* (×10^-3^)** | ***π* (×10^-3^)** | ***n*** | ***S*** | **Total bp** | ***θ_w_* (×10^-3^)** |
| **Chimpanzee** |  |  |  |  |  |  |  |  |  |  |  |  |  |  |
| Western | 9 | 231 | 2496576 | 0.034 | 0.027 | 7 | 276 | 211364 | 0.533 | 0.507 | 10 | 738892 | 160880685 | 1.623 |
| Eastern | 3 | 540 | 2496576 | 0.144 | 0.144 | 3 | 331 | 211364 | 1.044 | 1.044 | 12 | 702539 | 160880685 | 1.446 |
| Nigeria-Cameroon | 4 | 975 | 2496576 | 0.213 | 0.196 | 4 | 317 | 211364 | 0.818 | 0.809 | 20 | 401273 | 160880685 | 0.703 |
| **Bonobo** | 4 | 3284 | 3637523 | 0.492 | 0.459 | 4 | 220 | 268948 | 0.446 | 0.440 | 26 | 532077 | 160880685 | 0.867 |

^a^ Data from Hallast, et al. (2016); ^b^ Data from Nam, et al. (2015) and Chromosome 1 is used to represent autosomes.

*n*: sample size of chromosomes (rather than individuals); *S*: the number of segregating sites; *π*: nucleotide diversity calculated by Hallast, et al. (2016).

*θ_w_*: genetic diversity estimated by counting the number of segregating sites (i.e., Watterson estimator (Watterson 1975)).

Supplementary Figs. 1 to 5

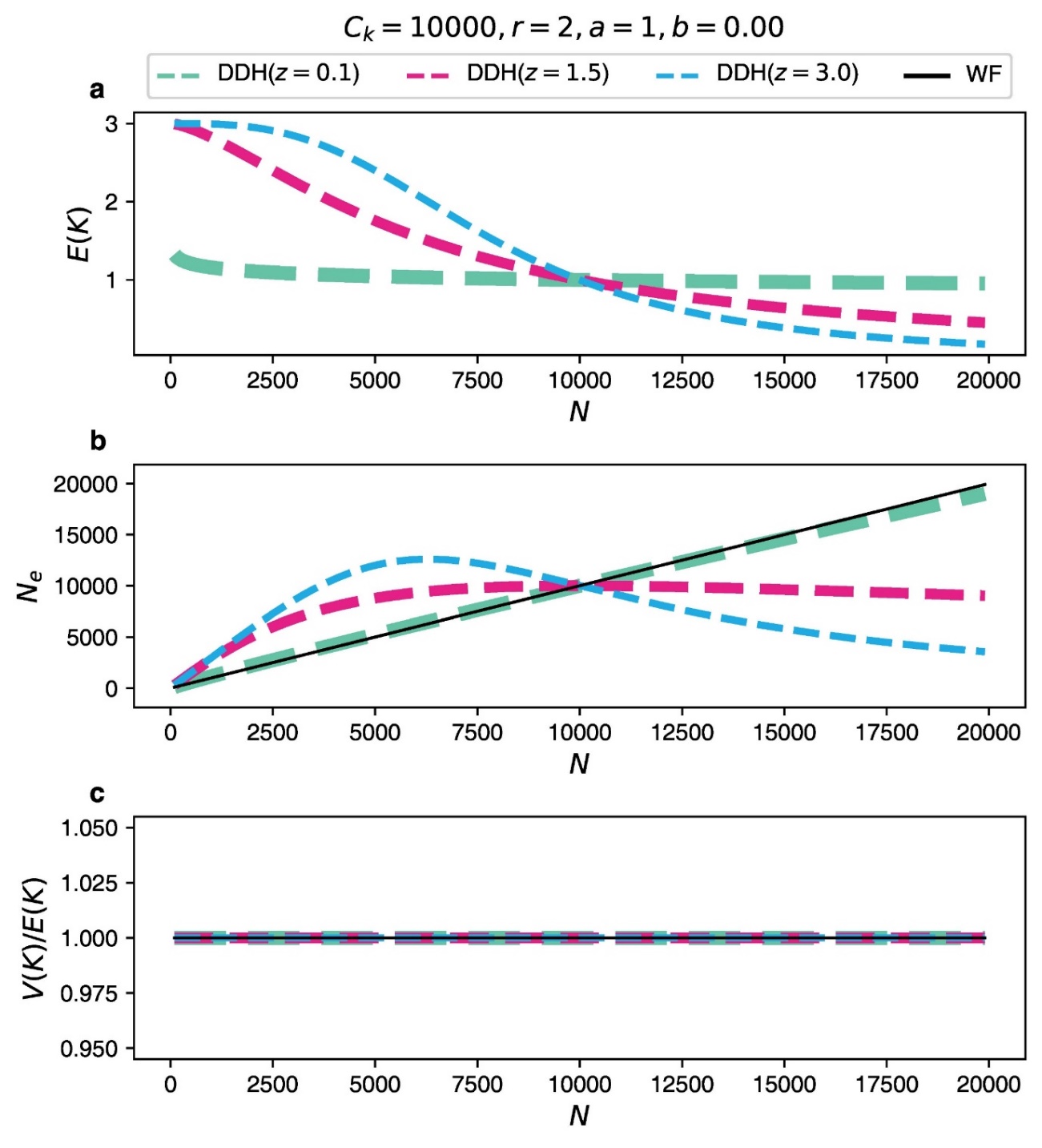

**Supplementary Fig. 1. Genetic drift under the density-dependent Haldane (DDH) model when the ratio of *V*(*K*) to *E*(*K*) keep constant as the population size increases.** Here we set the parameter *b* = 0.00 to let *V*(*K*)/*E*(*K*) ratio (= 1/*p_t_* – 1 = *a*(*N_t_*)*^b^*) be constant as the growth of the population (**c**). As the population size *N_t_* increases, the average number of offspring, *E*(*K*), decreases (**a**), resulting from the injuries number that an individual could tolerate decreases. However, the strength of genetic drift can indeed increase, decrease or stay nearly constant as population size *N_t_* changes. That’s to say, the effective population size *N_e_* could indeed decrease, increase or stay nearly constant as the growth of population size (**b**), depending on the ecology that governs *E*(*K*), *V*(*K*) and *N_t_*. The common parameters for the three panels are shown at the top of this figure.

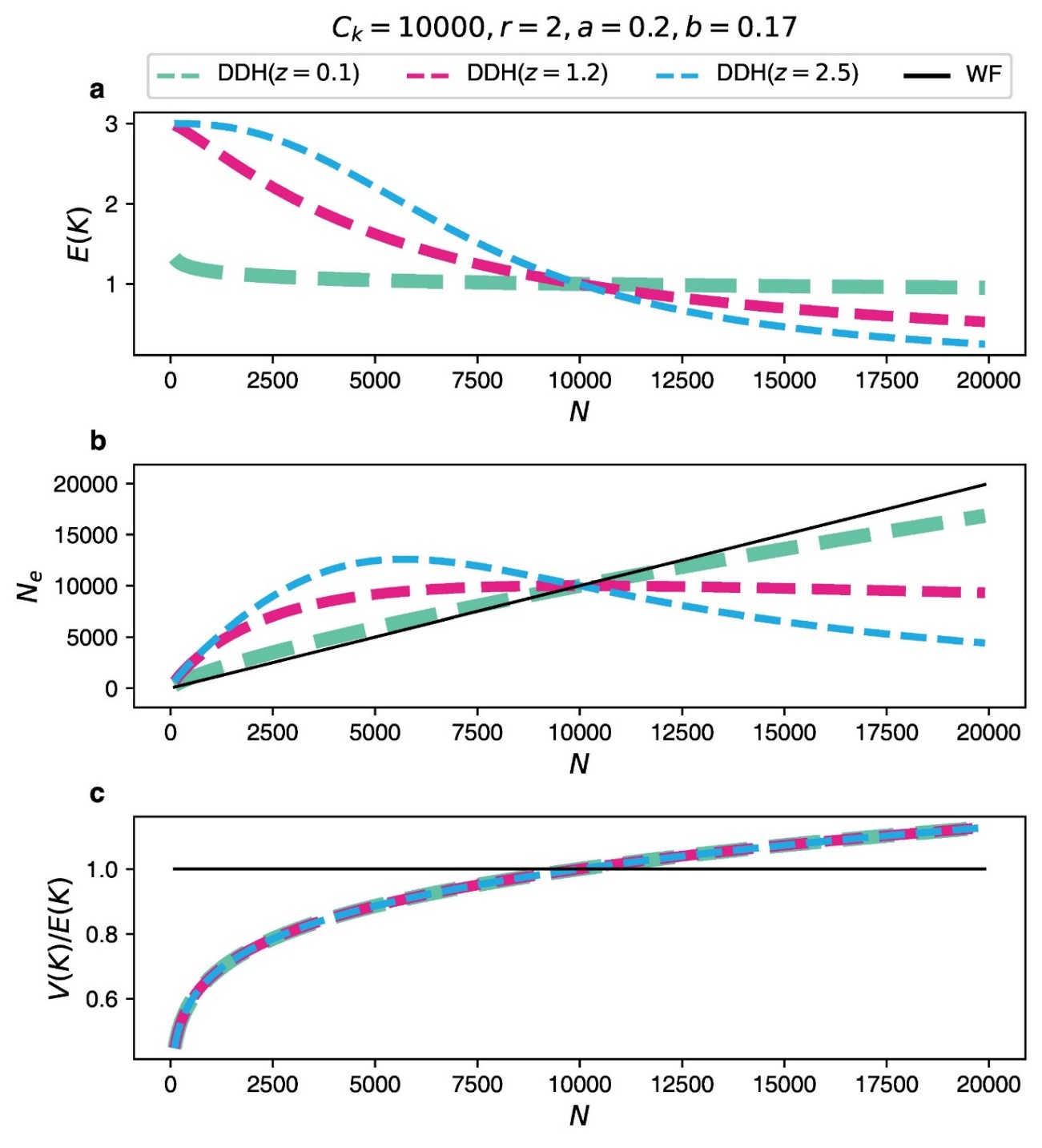

**Supplementary Fig. 2. Genetic drift under the density-dependent Haldane (DDH) model when the ratio of *V*(*K*) to *E*(*K*) increases as the population size increases.** Here we set the parameter *b* = 0.17 to let *V*(*K*)/*E*(*K*) ratio (= 1/*p_t_* – 1 = *a*(*N_t_*)*^b^*) increase as the growth of the population (**c**). As the population size *N_t_* increases, the average number of offspring, *E*(*K*), decreases (**a**), resulting from the injuries number that an individual could tolerate decreases. However, the strength of genetic drift can indeed increase, decrease or stay nearly constant as population size *N_t_* changes. That’s to say, the effective population size *N_e_* could indeed decrease, increase or stay nearly constant as the growth of population size (**b**), depending on the ecology that governs *E*(*K*), *V*(*K*) and *N_t_*. The common parameters for the three panels are shown at the top of this figure.

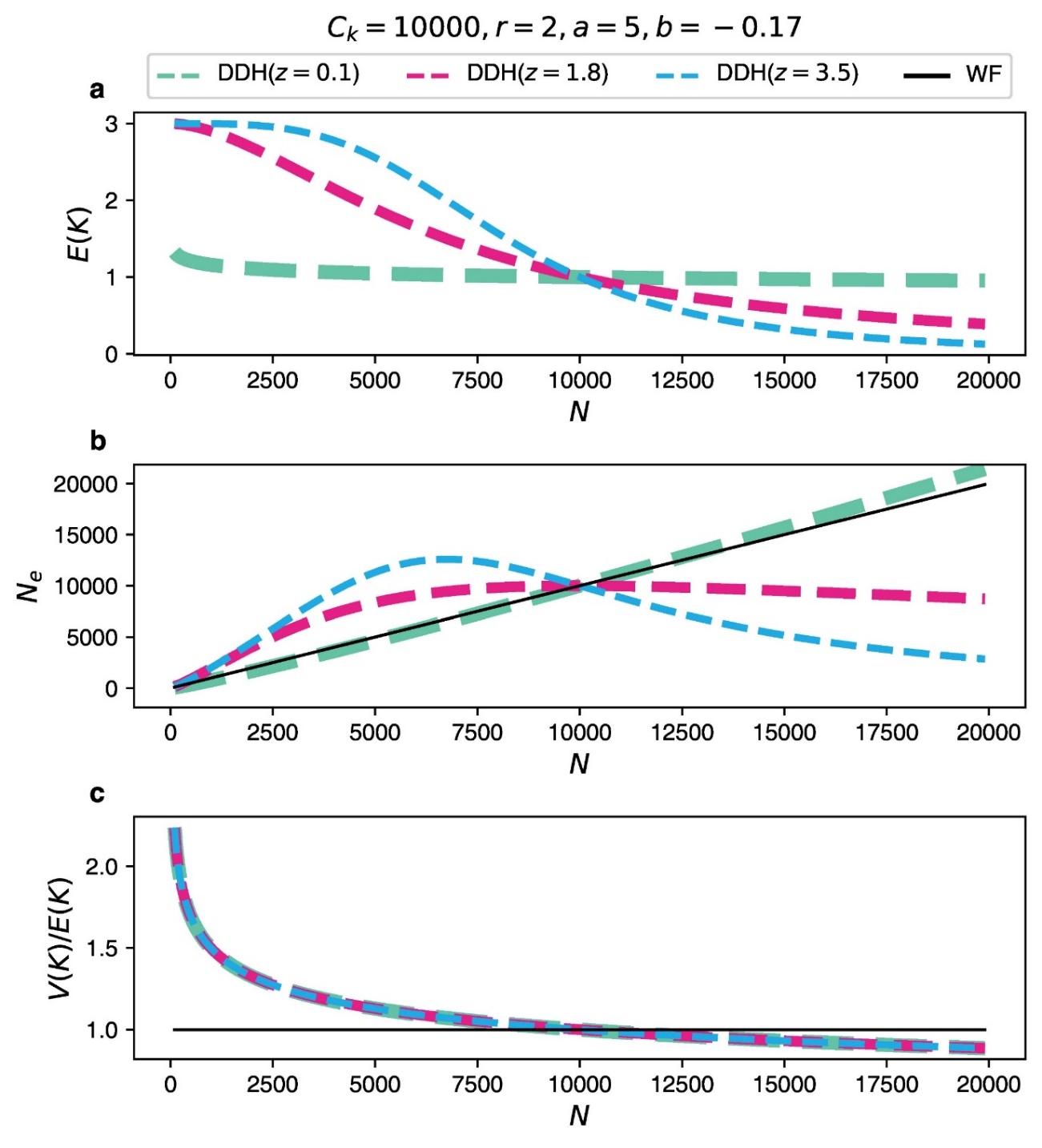

**Supplementary Fig. 3. Genetic drift under the density-dependent Haldane (DDH) model when the ratio of *V*(*K*) to *E*(*K*) decreases as the population size increases.** Here we set the parameter *b* = -0.17 to let *V*(*K*)/*E*(*K*) ratio (= 1/*p_t_* – 1 = *a*(*N_t_*)*^b^*) decrease as the growth of the population (**c**). As the population size *N_t_* increases, the average number of offspring, *E*(*K*), decreases (**a**), resulting from the injuries number that an individual could tolerate decreases. However, the strength of genetic drift can indeed increase, decrease or stay nearly constant as population size *N_t_* changes. That’s to say, the effective population size *N_e_* could indeed decrease, increase or stay nearly constant as the growth of population size (**b**), depending on the ecology that governs *E*(*K*), *V*(*K*) and *N_t_*. The common parameters for the three panels are shown at the top of this figure.

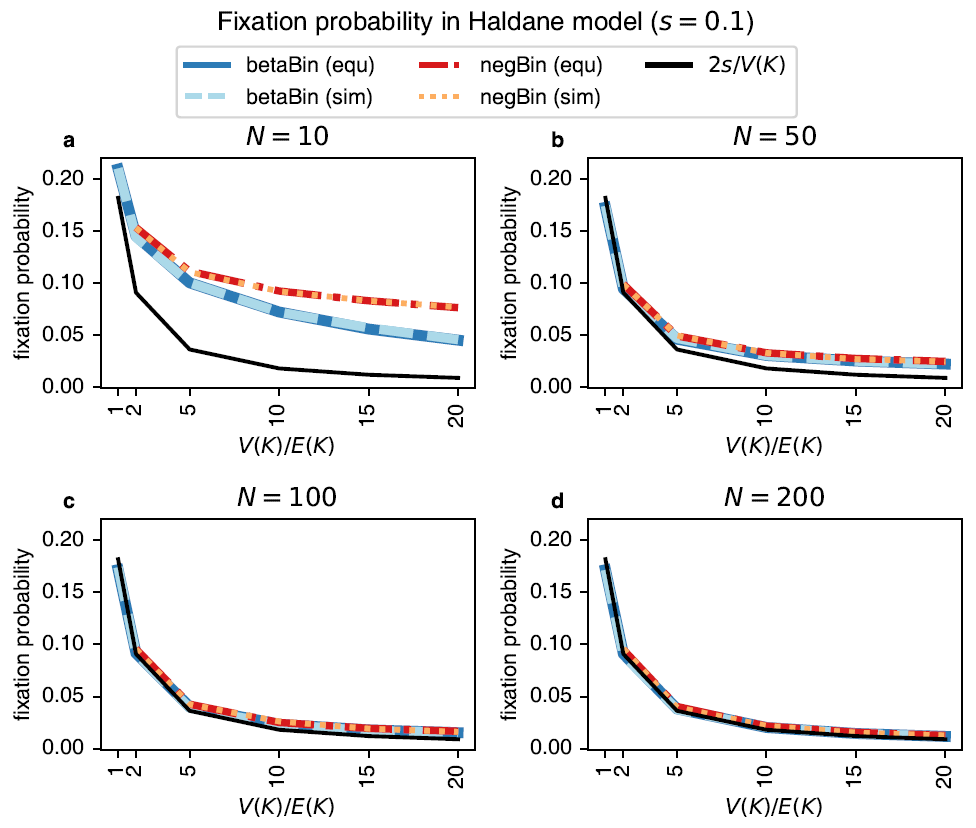

**Supplementary Fig. 4. Fixation probability of a new advantageous mutation in the Haldane model.** The fixation probabilities of advantageous mutations with the selective advantage of *s* = 0.1 are calculated based on approximate solution from Eq. (A3) as well as numerical solution from Eq. (A14) for the Haldane model. And their accuracies are confirmed by the simulation based on branching process. In all panels, the numerical solution from Eq. (14) is the same as the simulated value. **(a-b)** When *N* < 50, the approximate fixation probability from Eq. (A3) (the black line) is lower than the simulated values (the color lines) due to population extinction. **(c-d)** By the Haldane model, t the approximate fixation probability from Eq. (A3) is accurate when *N* reaches 100, which should be the case for most natural populations. Details: In the Haldane model, the number of progenies for the mutant and wildtype is denoted as *K_M_* and *K_W_*, respectively. We let *E*(*K_W_*) = 1, *E*(*K_M_*) = 1 + *s.* *V*(*K*)/*E*(*K*) is the same for the two alleles. In the Haldane model, both *K_M_* and *K_W_* follow the beta-binomial (betaBin, blue lines) or negative binomial (negBin, red lines) distribution. In each of the four panels**,** the fixation probability is shown as a function of *V*(*K*)/*E*(*K*). The black line shows the approximate fixation probability of 2*s*/*V*(*K_M_*).

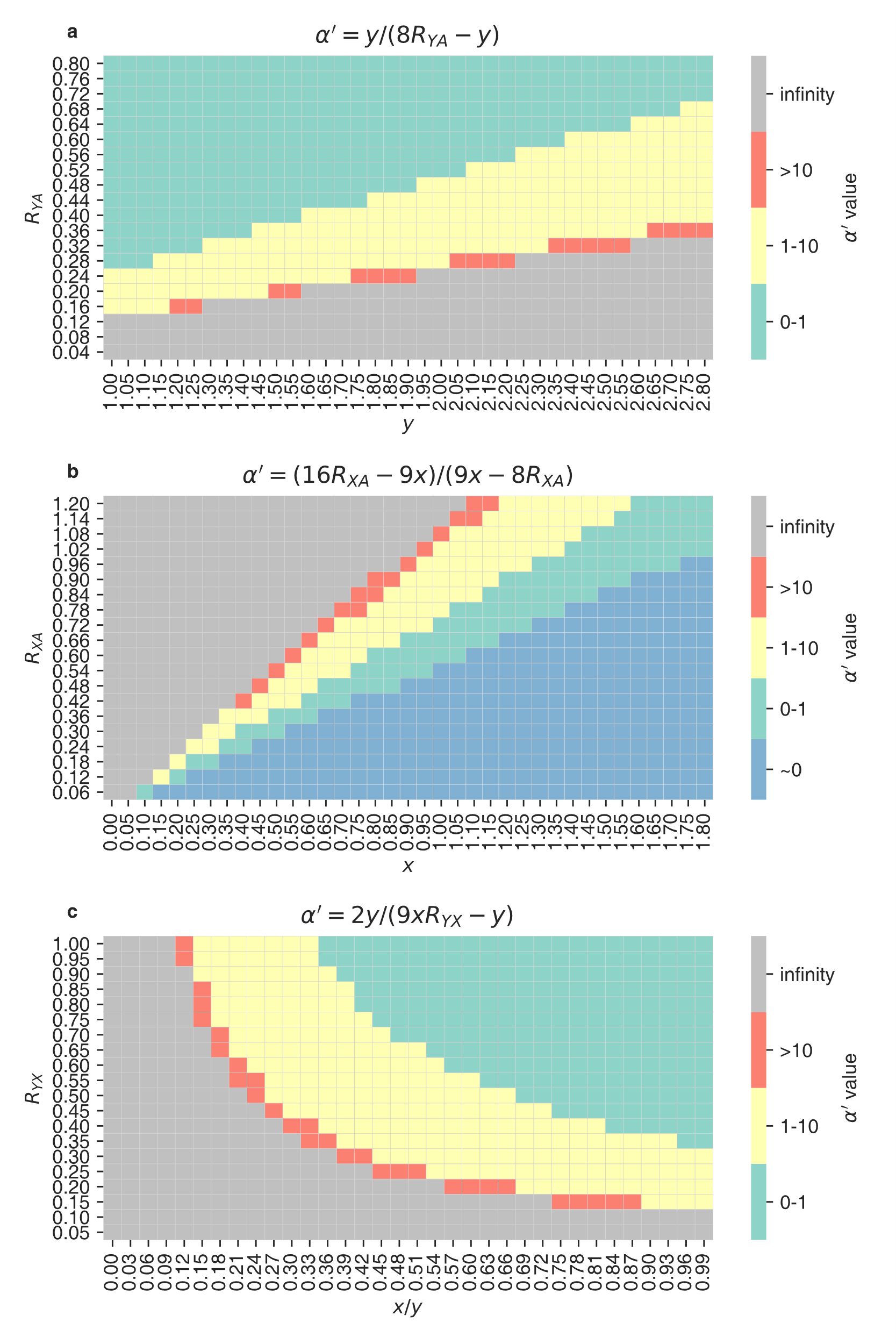

**Supplementary Fig. 5. The sensitivity of estimation for *α'*.** *y* and *x* are the mutation rate ratios for Y/A and X/A. *R_YX_* (=*θ_Y_* /*θ_X_*), *R_YA_* (=*θ_Y_* /*θ_A_*) and *R_XA_* (=*θ_X_* /*θ_A_*) are the Watterson genetic diversity ratios for Y/X, Y/A, and X/A. The *α'* value could be estimated by three methods: (**a**) a function of *R_YA_* and *y* from Eq. (A35.1); (**b**) a function of *R_XA_* and *x* from Eq. (A35.2). Note the negative estimated value for *α'_XA_* has two possible meaning due to non-monotonic function of *x* and *R_XA_*: *α'* is infinity (i.e., *V_m_* ≫ *V_f_*) when *α'_XA_* ≤ -2 or *α'* is approximate to zero (i.e., *V_m_*/*V_f_* ~ 0) when -1 ≤ *α'_XA_* < 0; (**c**) a function of *R_YX_*, *x*/*y* from Eq. (A35.3). Note the narrow red band where *α'* > 10.
